# Simultaneous photoactivation and high-speed structural tracking reveal diffusion-dominated motion in the endoplasmic reticulum

**DOI:** 10.1101/2023.04.23.537908

**Authors:** Matteo Dora, Christopher J. Obara, Tim Abel, Jennifer Lippincott-Schwarz, David Holcman

## Abstract

The endoplasmic reticulum (ER) is a structurally complex, membrane-enclosed compartment that stretches from the nuclear envelope to the extreme periphery of eukaryotic cells. The organelle is crucial for numerous distinct cellular processes, but how these processes are spatially regulated within the structure is unclear. Traditional imaging-based approaches to understanding protein dynamics within the organelle are limited by the convoluted structure and rapid movement of molecular components. Here, we introduce a combinatorial imaging and machine learning-assisted image analysis approach to track the motion of photoactivated proteins within the ER of live cells. We find that simultaneous knowledge of the underlying ER structure is required to accurately analyze fluorescently-tagged protein redistribution, and after appropriate structural calibration we see all proteins assayed show signatures of Brownian diffusion-dominated motion over micron spatial scales. Remarkably, we find that in some cells the ER structure can be explored in a highly asymmetric manner, likely as a result of uneven connectivity within the organelle. This remains true independently of the size or folding state of the fluorescently-tagged molecules, suggesting a potential role for ER connectivity in driving spatially regulated biology in eukaryotes.

## I. INTRODUCTION

One of the most notable aspects of eukaryotic life is the development of membrane-enclosed organelles [6]. These compartments provide two important advantages to cells. First, they segregate unique chemical environments that allow incompatible biochemical reactions to occur simultaneously, and second, they provide extensive two-dimensional surfaces (membranes) to facilitate the efficiency of biological reactions [2]. The largest organelle in eukaryotes is the endoplasmic reticulum (ER). Topologically, the ER is a single, immense compartment that divides the nucleoplasm from the cytoplasm and stretches to the furthest reaches of the cell as a structurally complex membrane-enclosed network [25]. It is a major site of cellular translation, protein folding, and quality control [60, 62], the major cellular calcium store and sink in most cells [8], and the primary site of lipid synthesis and regulation [77]. Thus, understanding how molecules are spatially distributed over time within the ER network is important for understanding how their functions are achieved.

The development of technologies for fluorescent tagging of molecules has opened the possibility to probe the dynamic nature of proteins within the ER [17, 75]. In particular, fluorescence recovery after photobleaching (FRAP) and fluorescence lifetime in photobleaching (FLIP) [39] have provided significant insight into the way many components of the compartment behave [41]. These studies have suggested that the composition of the organelle is strikingly dynamic, equilibrating on a timescale of seconds [11, 50]. However, these studies are limited to single structurally heterogeneous spatial regions (FRAP) or time scales of tens of seconds (FLIP), and the ability to simultaneously observe high-speed ensemble dynamics across the entire cell has proven elusive. The development of photoactivatable fluorophores [57] has enabled this in other systems [30, 52], but most photoactivatable fluorophores do not fold correctly or provide low SNR in the high redox environment of the ER lumen [12, 33].

An additional limitation to understanding protein redistribution within the ER is provided by the convoluted structure of the organelle itself [73]. The small size of ER substructures necessitates that molecules are not free to move uniformly in any dimension. Previous work has attempted to account for this in FRAP data with a number of clever modeling-based approaches, but the inability to directly observe the structure in question during fluorescent recovery has led to inconsistent results from varying model assumptions [55, 66, 67, 71].

In this article, we develop a combined experimental and analysis pipeline to surmount these two issues. We have used photoactivation of protein-linked organic dyes that work well within the ER lumen to track the dynamic distribution of ER-localized molecules. During photoactivation, we simultaneously image the underlying structure of the organelle with an independent camera and orthogonal fluorescent label. This is approach supports high speed imaging while also collecting entire cells, and allows us to place the photoactivated material back in the structure it is localized to in real time. We then coupled this to a novel, machine learning-assisted analysis pipeline to provide a first-order approximation of the complex organelle structure, allowing estimation of the potential paths taken by the fluorescent components. Collectively, this pipeline allows the extraction of quantitative information about the molecular behaviour within the ER at the whole-cell scale.

## II. RESULTS

In order to track ER protein dynamics across entire cells, we utilized photoactivatable organic dyes that can be directly coupled to a protein of interest by genetic fusion to a HaloTag [18, 26, 42]. These dyes are functional in the ER lumen and are not thought to induce the misfolding and ER stress-related pathways common with fluorescent proteins not optimized for the high redox environment within this compartment [13]. Once labelled with the dye, proteins of interest can be fluorescently marked at specific locations within the ER structure by stimulation with 405nm light, and their distribution over time can be monitored by high speed microscopy. We performed experiments in cells also expressing a non-perturbative GFP-derivative (moxGFP) targeted to the ER lumen [12], to provide a simultaneously collected map of the underlying structure of the ER for downstream analysis purposes.

Briefly, COS-7 cells were cotransfected with moxGFP and a HaloTag fused to the protein construct of interest. We then used high speed spinning disk microscopy (at 10 Hz acquisition rate) using dual cameras to simultaneously collect the distribution of the photoactivated species of the dye and the fluorescence of the GFP label. This resulted in two stacks of paired 2D images collected at continuous 100 ms intervals. Photoactivation was carried out continuously at single, diffraction limited spot, and the diffusion of molecules away from that spot was visible as the distribution of the converted dye molecule (fig 1).

**Figure 1.**
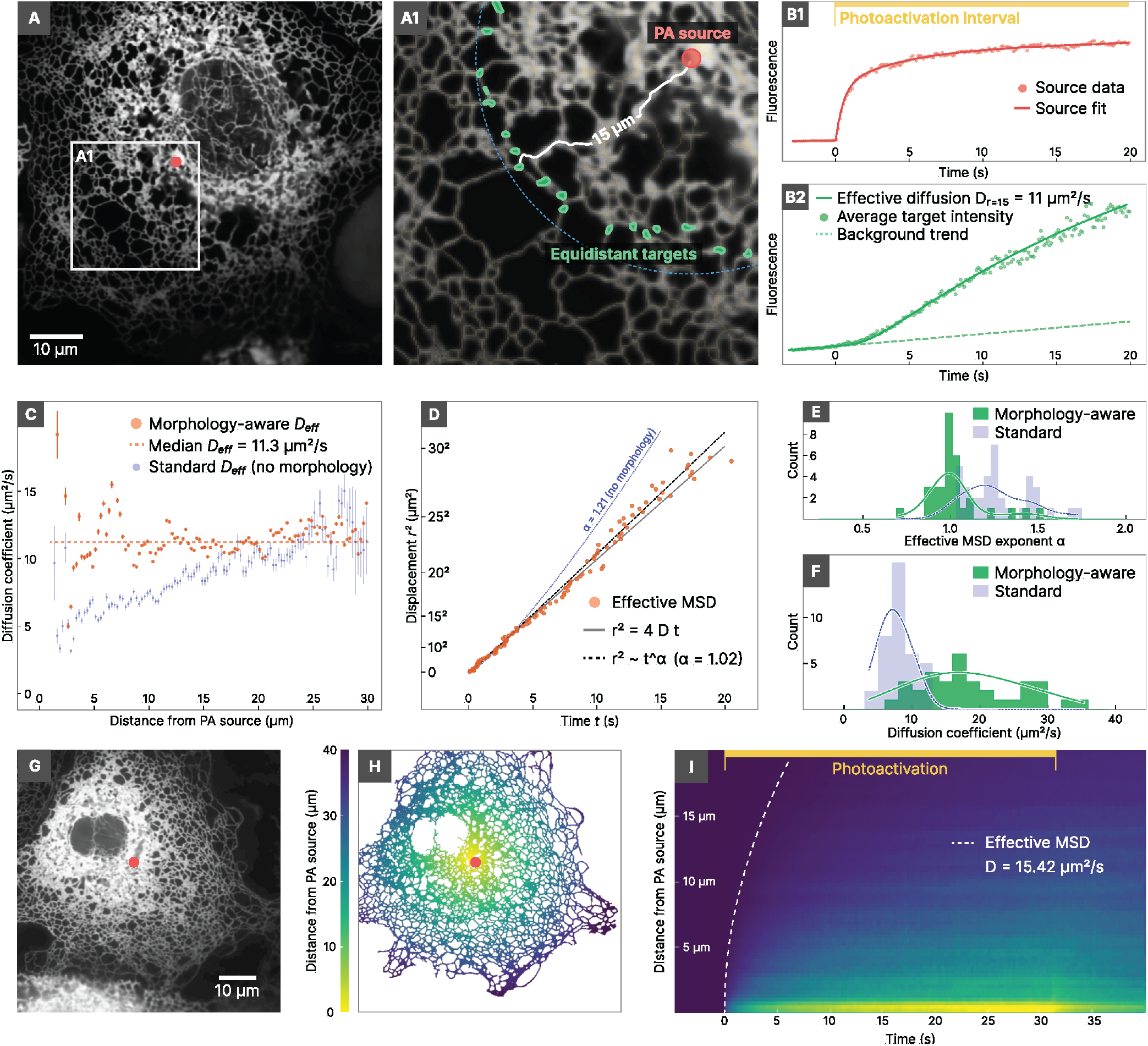
Brownian diffusion governs large scale ER luminal dynamics. **A** The ER structure of a COS-7 cell. The red dot indicates the location of photoactivation. Green regions correspond to ER domains that are equidistant (d = 15 μm) from the photoactivation region (distance calculated on the underlying ER morphology). The blue dashed line denotes the Euclidean distance of 15μm from the photoactivation region when the ER morphology is neglected. **B1** Data and analytical fit of the fluorescence curve inside the photoactivation region. **B2** Average fluorescence of the target regions at distance 15 μm from the photoactivation source and t of effective diffusion model (D = 11 μm^2^ s^−1^). **C** Effective diffusion coefficient as a function of the distance *d* from the photoactivation region, obtained from separate fits at each distance, for the morphology-aware model (orange) and the standard model neglecting ER morphology (light blue).The error bars represent the approximated confidence interval estimated by observed Fisher information (± 5 standard deviations). **D** Effective mean-squared displacement (MSD) obtained by ensemble estimate. **E** Distribution of the effective MSD exponent *α* of each cell for the morphology-aware and the standard model (N = 33 and N = 37 respectively), obtained by ensemble estimate. **F** Distribution of the median cellular diffusion coefficients obtained by ensemble estimate for HaloTag-KDEL in the ER lumen, comparing the morphology-aware model (green) and the standard model neglecting morphology (light blue). **G** Example of ER in a COS-7 cell. The photoactivation region is denoted by the red dot. **H** Structure of the ER network of the cell in G with colour scale denoting distances from the photoactivation source. **I** Kymograph of the average fluorescence in time of the cell in G, binned over the distance from the photoactivation source through the structure. The white dashed line denotes the squared root of effective MSD curve of the diffusion model that was fit 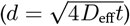.

### A. Ensemble diffusion governs dynamics of ER luminal content

We first set out to evaluate the dynamics of lumen-resident proteins that are freely floating (i.e. without membrane anchors or known specific binding partners). We assumed a globally isotropic but not necessarily diffusive motion, since local directional flows may be present [31]. To accomplish this, we used a HaloTag targeted to the ER lumen with a signal sequence and a c-terminal KDEL retention motif as a model probe for free-oating luminal content (HaloTag-KDEL).

Proteins within the ER lumen are constrained to compartment by the bounding membrane, so they are not free to move in any arbitrary direction. Thus, analysis of the distribution patterns must account for the underlying organelle structure, or model fitting will be prone to artefacts (e.g., [73]). To address this, the ER was first segmented by analysing the signal from the moxGFP to extract the network structure (see *Methods*, section IV A and supplementary figure S1). Regions of the ER were characterized by their distance from the PA source as measured through the underlying structure of ER (fig. 1A, inset), as opposed to Euclidean distance (fig. 1A, dashed line).

Using this approach, the estimated arrival time of fluorescent molecules can be coupled with the distance from the PA source and the amount of photoactivation to produce an effective diffusion model. In principle, this model depends on the rate of photoconversion by the photoactivation laser, which cannot be easily determined. To circumvent this issue, we take a hybrid data-driven modelling approach. We first quantified the photoactivated fluorescence over time at the PA source and fitted it with an arbitrary parametric function *ϕ* (*t*). In our case, we found that a function *ϕ* (*t*) in the form of a sum of saturating exponentials could well describe the data (fig 1B1, and see *Methods*). We then fixed the effective diffusion model conditioning it on *ϕ* (*t*) at the source region, effectively constraining the model based on the measured intensity of the PA source. Once the source parameters were determined, we performed a maximum-likelihood the of the fluorescence intensity model to extract the effective diusion coefficients corresponding to the average fluorescence measured in the equidistant points (fig 1B2). We constrained the model parameters based on the density of the ER structure, which we estimated directly from the moxGFP channel (*Methods*, section IV C). e fluorescence intensity model also considers the possible background photoactivation (fig 1B2, dashed line) due to inefficient photoactivation of the dye by the laser used to image the GFP. Note that this approach can be modified to deal with sources of anisotropy if the model t is not good, as would be predicted by synchronous directed flows or other micron scale sources of anisotropy (see section E below). We then repeated this procedure for every distance value in the structure, obtaining distance-dependent estimations of the effective diffusion coefficient. In fig 1C we compare, on an example cell, the results of this morphology-aware model (orange) to a standard maximum-likelihood fit that only considers Euclidean distances and that does not impose constraints based on ER density (light blue). We note that the approach that does not take ER morphology into account produces estimates of the diffusion coefficient that increase with the distance from the PA source (an indication of super-diffusive motion), while such effect disappears when considering a morphology-aware model.

In the immediate proximity of the PA source both approaches are highly sensitive to variations in fluorescence and can badly estimate the effective diffusion coefficient. Similarly, the quality of the t degrades in regions that are too far away from the PA source, since only a relatively small fraction of molecules can reach those regions over the considered timescale, thus decreasing the signal-to-noise ratio of the measured fluorescence. To mitigate these issues, we estimate the uncertainty of the diffusion coefficient fit based on observed Fisher information (see *Methods*, section IV C), discarding data points with standard deviation above a threshold of 0.1 μ m^2^ s^−1^.

To evaluate the validity of a diffusion model more generally, we represented the fit results for each cell in the form of effective diffusion timescales, creating an effective MSD curve describing the time evolution of the average distance travelled by a protein released from the PA source (fig. 1D). The exponent α of the MSD curve allowed us to evaluate possible deviations from standard diffusion (α = 1), such as sub diffusion (α < 1) or super diffusion (α > 1). For each cell, we determined the MSD exponent α by fitting the effective MSD data with a function of the form (c t^α^) (fig. 1D), comparing the proposed morphology-aware model (black dashed line) with the MSD obtained from a standard model that neglects ER morphology (blue dotted line). We found that population statistics of cells expressing HaloTag-KDEL had an average MSD exponent α ≈ 1, thus suggesting that luminal dynamics can be well described by standard diffusion over this spatiotemporal scale (fig. 1E, green). We also note how neglecting ER morphology resulted in apparent super-diffusive behaviour with α > 1 (fig. 1E, light blue). This is a surprising contrast to modeling performed in the peripheral ER, where erroneous model fits as a result of structural confinement generally manifest as subdiffusive scaling of the MSD exponent (likely the result of lower Renyi dimensionality and competing molecular populations, see *Discussion* and *Supplementary Text and Discussion*, Sections 1 and 5) [55, 66, 67, 73]. However, in agreement with these studies, model fits that account for the structure of the ER show significantly faster diffusion rates than those that neglect them (fig. 1F).

For organelles with less complex morphology, changes in the distribution of fluorescent molecules over time are often effectively represented with a kymograph, i.e. a time-space plot of the fluorescence observed along one dimension over time [32, 45, 85]. In the ER, this is challenging due to the circuitous and dynamic nature of paths through the structure and the relatively low number of fluorescent molecules present in any isolated structural component. As a visualization tool, we used the distances calculated between the PA source and every point of the segmented ER structure (fig. 1G–H), andaveraged the fluorescence values at each distance to create an equivalent kymograph describing the time-evolution of the fluorescence as a function of the distance from the PA source (fig. 1I). This representation shows concentrated photoactivated signal at the bottom (close to the PA source) and a gradual spreading through the structure moving upwards (towards distant regions) as the time of photoactivation increases. As a qualitative evaluation of the diffusion model fit to this cell, the time evolution of the fluorescence is continuous and well contained within the bounds predicted by pure diffusion at the effective diffusion coefficient observed, both notable differences from what we have theoretically predicted for active flows on visible spatial and temporal scales [16].

Specifically, one prediction of our previous work [16, 31] is that active flows in luminal content are predicted to form fluorescent “packets” when they occur on observable scales. Although we did not observe such packets in the spatially averaged kymographs (fig. 1I), we wondered whether the large degree of complexity in the ER may be masking such phenomena based on heterogeneity of packet distances from the PA source. To address this, we chose an example cell that showed significant local heterogeneity in the photoactivated channel (fig. 2A) and simulated purely diffusive motion directly on the experimentally-obtained ER morphology. Briefly, the proposed effective diffusion model was implemented with constant diffusion coefficient defined as the median of the maximum-likelihood fit for the cell (see *Methods*, section IV C).

**Figure 2.**
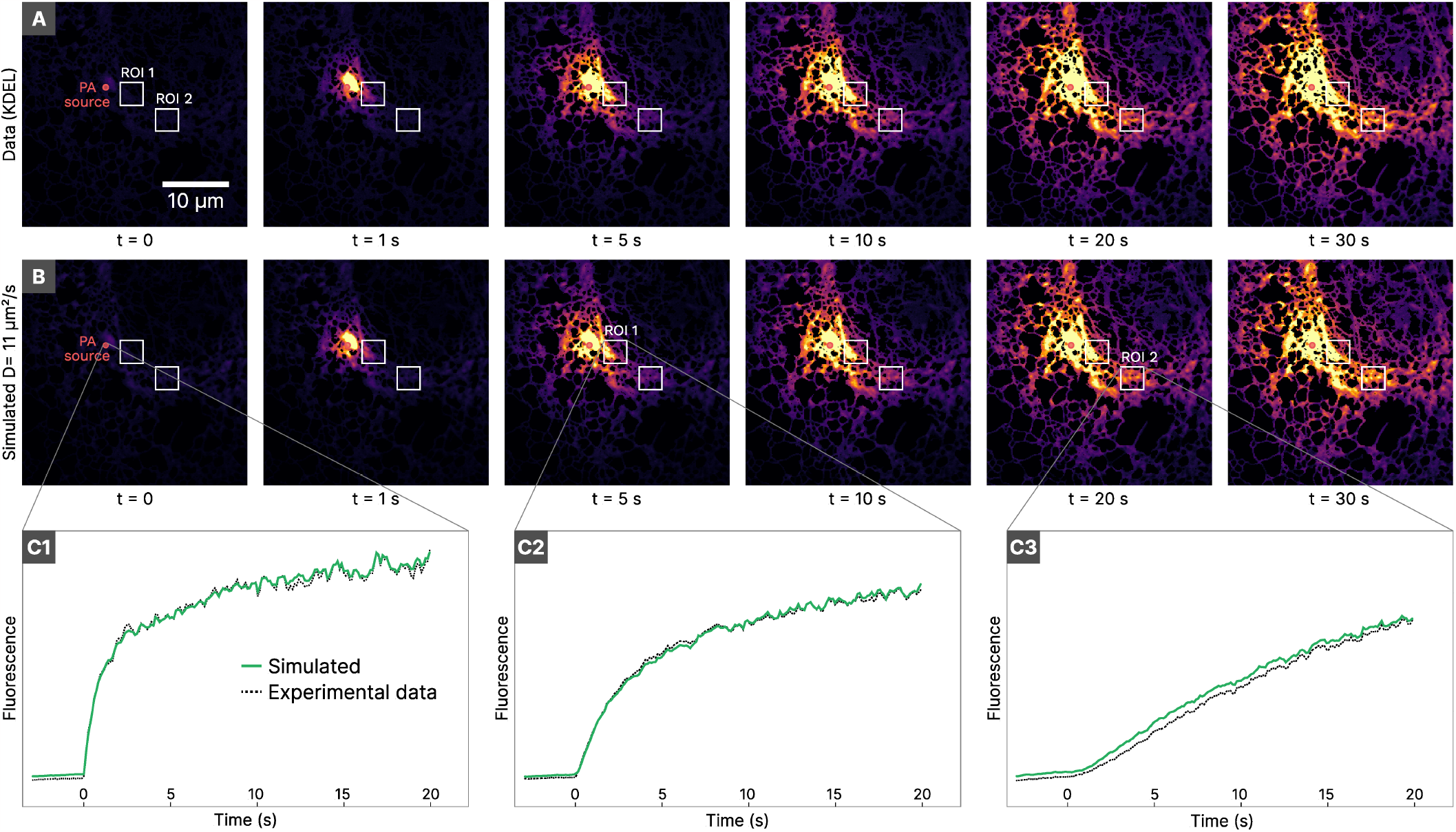
**A** Experimental snapshots at different time intervals from the beginning of photoactivation. **B** Simulated snapshots built by evaluating the effective diffusion model on the ER shape obtained by experimental data, using the median diffusion coefficient obtained by the morphology-aware maximum-likelihood fit. **C** Comparison of experimental and simulated fluorescence curves at the photoactivation source (C1) and at two ROIs (C2 and C3) located at different distances from the PA source.

The data was then graphically applied to the underlying ER structure, weighted by the underlying moxGFP fluorescence (fig. 2B, see *Supplementary Text and Discussion*, Section 2). Remarkably, the simulated distribution was nearly indistinguishable from the experimentally-observed photoactivation channel, correctly predicting fluorescence arrival times in both bright and dim regions of the structure (fig. 2C). Notably, even the fluctuations in fluorescence signal were largely predicted by the model, suggesting that the majority of the heterogeneity observed in the photoactivated channel is due to the varying density and motion of the underlying ER structure and not to uneven mixing of the molecules.

### B. Effects of membrane anchoring on ER protein diffusion

Classic work in the field using numerous methods has established that viscous drag on transmembrane domains is a dominating factor on the diffusive properties of membrane-associated proteins (reviewed in [14]). However, the degree to which this drag slows motion compared to freely diffusing molecules has been inconsistently reported, likely as a result of varying model assumptions and tools [22, 58, 66, 71]. Since our approach uses data to directly estimate the parameters that are taken as boundary conditions in these previous studies, we reexamined the phenomenon with knowledge of the underlying structure and dynamics of the ER.

To achieve this, the freely floating HaloTag in the ER lumen (HaloTag-KDEL, introduced in fig. 1) was compared to the same HaloTag anchored to the ER membrane. The membrane anchoring was achieved by fusing HaloTag genetically to a targeting domain from Sec61*β* that is known not to interact with the translocon in living cells [53]. Photoactivation experiments were then carried out exactly as described above, monitoring the arrival times of photoactivated HaloTag-Sec61 *β* throughout the structure of the ER.

In agreement with the literature, membrane-anchored HaloTag showed significantly reduced distances traveled over the same time window, compared to HaloTag free in the ER lumen (fig. 3A-B). This resulted in a smaller fraction of the cell falling within the quality threshold for analysis, but the relatively larger fraction of molecules within that space caused significantly more uniform and stable signal in each cell (fig. 3C). Estimation of the isotropic diffusion characteristics was consistent with published literature [35, 73] and the previous results with HaloTag-KDEL (fig. 3C), showing that membrane dynamics were also well described by standard Brownian diffusion (MSD exponent α = 1.02, fig 3D). Population level statistics for the effective diffusion coefficient (fig. 3E) confirmed that membrane proteins are diffusing significantly slower than luminal proteins in our system (membrane D_Sec61_*β* = 5.39 μm^2^ s^−1^, compared to luminal D_KDEL_ = 16.84 μm^2^ s^−1^, p-value < 1 × 10^−3^). Note that as with HaloTag diffusion within the lumen, accounting for the micron-scale ER structure results in estimates of ER membrane protein diffusion that are significantly elevated compared to literature values. This is consistent with our published work using single molecule tracking to look at HaloTag-Sec61*β* in the ER periphery, which also suggested morphology-unaware approaches dramatically underestimate the true diffusion of membrane proteins in convoluted structures like the ER [73] (see *Discussion* and *Supplementary Text and Discussion*, Section 3).

**Figure 3.**
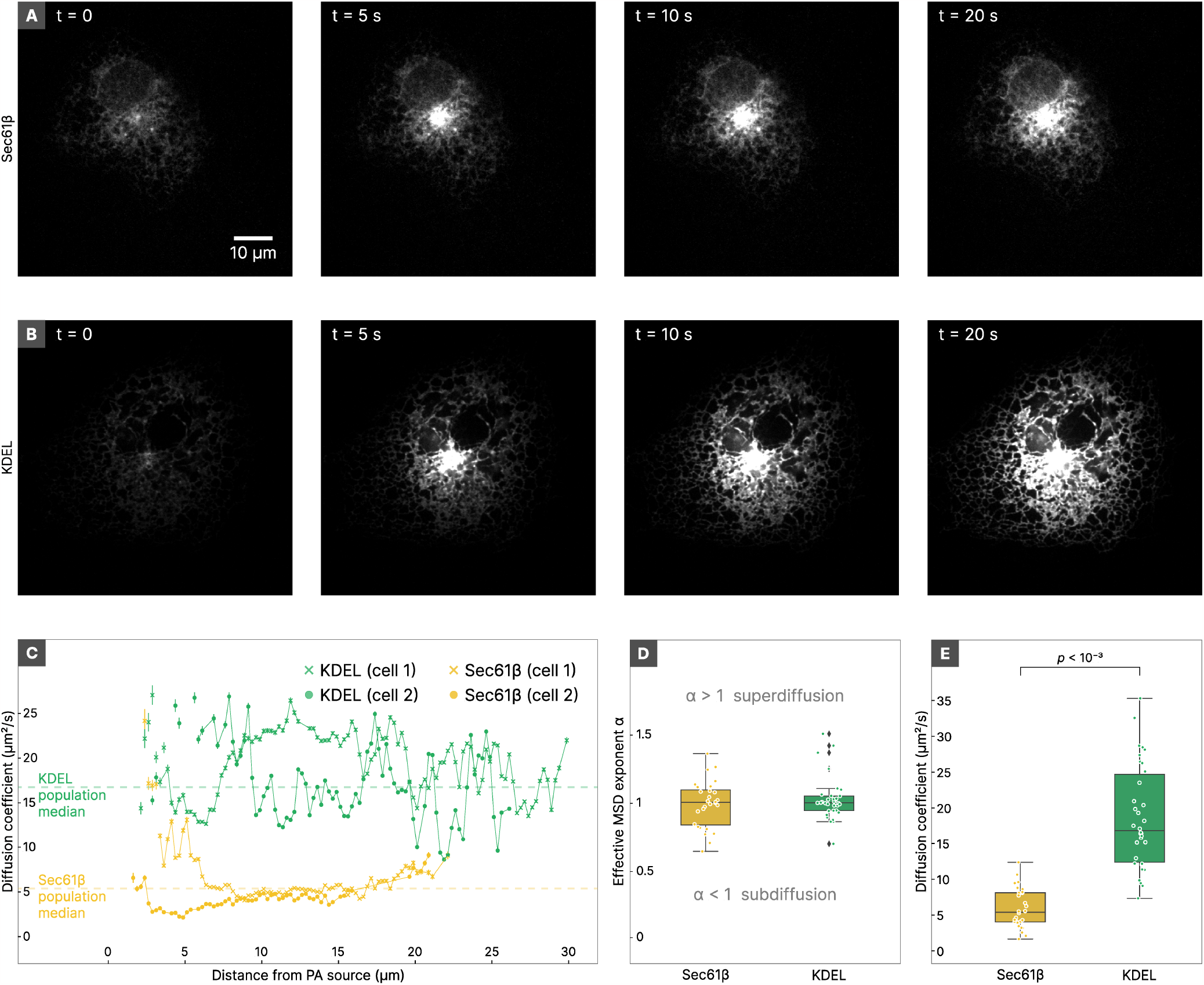
Comparison of ER protein dynamics in the lumen and in the membrane. **A-B** Examples showing the spatial distribution of photoactivated HaloTag at various time points post-activation when anchored to the membrane (Sec61,*α*)(A) or freely diffusing in the lumen (KDEL)(B). **C** Two representative examples of the isotropic diffusion t in a cell (averaged over all targets in the cell at a fixed distance) for HaloTag-KDEL and HaloTag-Sec61, *β*, plotted over the median value for all cells observed. Note the increased heterogeneity of effective diffusion coefficients for luminal content. **D** Cell population statistics of the effective MSD exponent for HaloTag when linked to Sec61, *β* or free in the ER lumen (KDEL). **E** Population statistics of the effective diffusion coefficient (*D*_eff_) of HaloTag-KDEL and HaloTag-Sec61, *β* conditions.

### C. Effect of protein size on mixing dynamics

Transmembrane proteins show significantly reduced diffusion as a result of viscous drag in the membrane, but classical FRAP and fluorescence correlation spectroscopy (FCS) experiments suggest that even the ER lumen is itself significantly more viscous than the cytoplasm (reviewed in [40]).The Stokes– Einstein formula predicts that the mean diffusion coefficient of freely diffusing molecules should be inversely proportional to the radius of the molecule. We tested our ability to resolve this relationship by generating a larger version of a free-floating molecule in the ER lumen. Briefly, we genetically fused a signal sequence to three codon optimized concatemers of the HaloTag linked by flexible linkers and C-terminally fused to a KDEL retention sequence (3×HaloTag-KDEL).

As predicted for a diffusion-dominated system, the photoactivated 3×HaloTag-KDEL explored a slightly smaller proportion of the cell in a defined time window than the single HaloTag-KDEL construct (fig 4A-B). Accordingly, it also showed much more stable mean arrival times over distance, similar to what was seen when HaloTag was slowed by a membrane anchor (fig 4C), suggesting this effect is a result of slower motion leading to more dense sampling in defined time windows and not inherent differences in the properties of the lumen as opposed to the membrane. Again, we found that the dispersion of the 3×HaloTag was most effectively fit with a model for simple Brownian motion (effective MSD exponent α_3×_=1.01, fig 4D). In agreement with physical characteristics of diffusion, we observed significantly decreased mixing speed for the 3×HaloTag-KDEL compared to single HaloTag-KDEL (D_3×_ = 8.53 μm^2^ s^−1^, D_1×_ = 16.84 μm^2^ s^−1^, p-value < 1 × 10^−3^, see fig 4E). Notably, as a control, the same 3×HaloTag fused to the membrane targeting domain of Sec61α had no statistically significant effect on its diffusion characteristics, since the viscous drag of the membrane on the membrane anchor was dominating and the 3x tag faced the cytoplasm (fig 4E). In agreement with this, we note that the mean effective diffusion coefficient of 3xHaloTag-KDEL is still nearly double that of the single HaloTag when it is anchored in the membrane (HaloTagSec61α, fig 4E).Thus, in agreement with other approaches in the literature, viscosity in the ER lumen is significant and can be detected with this method, but is still qualitatively dominated by the drag of membrane anchors.

**Figure 4.**
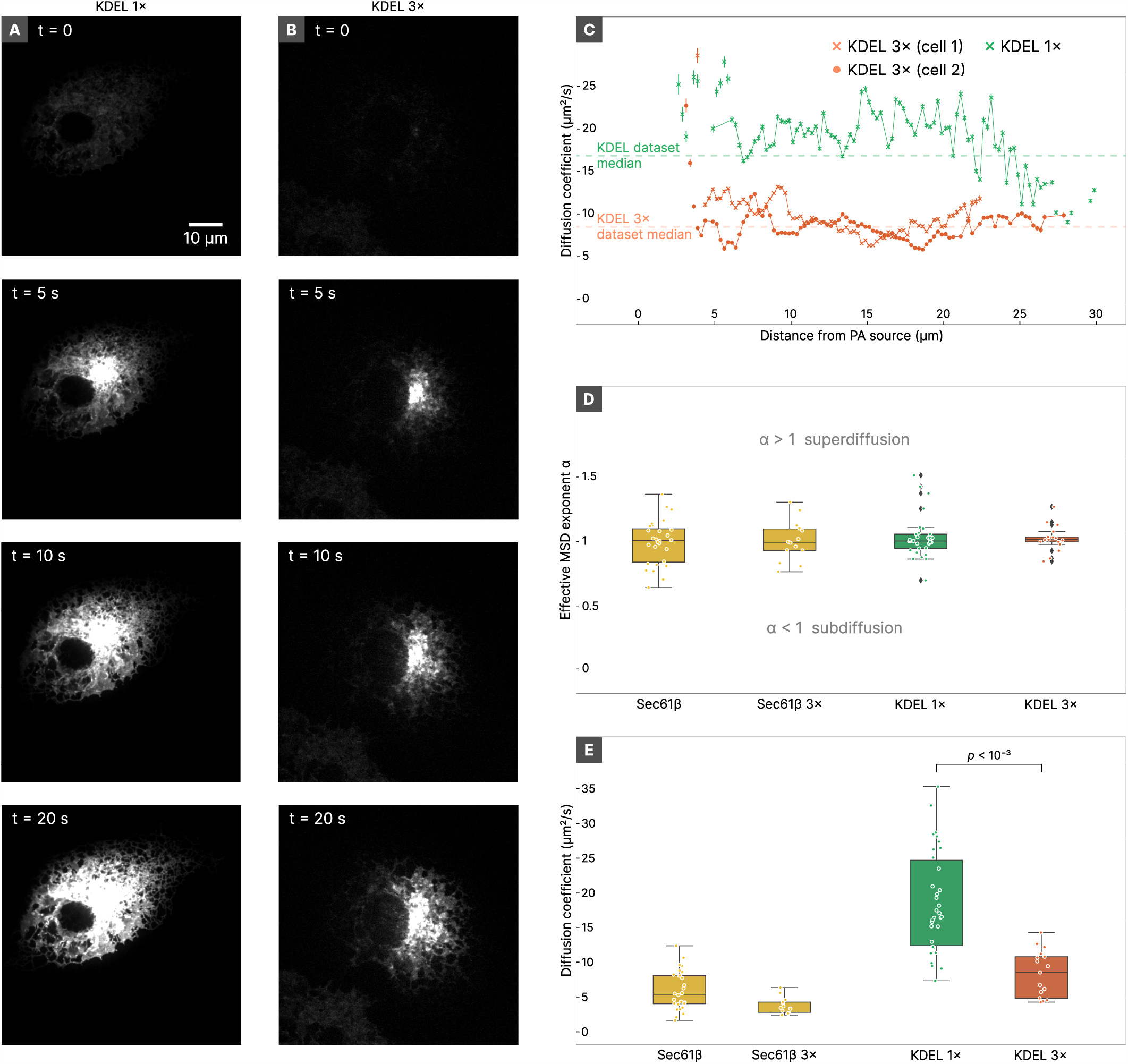
Diffusion in the ER lumen is protein size-dependent. **A-B** Examples of the spatial distribution of photoactivated HaloTagKDEL(A) or 3×HaloTag-KDEL(B) at defined time points after the start of photoactivation. **C** Results for the isotropic diffusion fit (averaged over all targets at fixed distance) for two example cells expressing 3×HaloTag-KDEL, compared to an example of HaloTag-KDEL. Note the reduced variance in effective diffusion coefficients with the larger luminal protein. All results are plotted over lines indicating the median value of all cells in the dataset. **D** Population statistics of the effective MSD exponent of for HaloTag-Sec61, *α*, 3×HaloTag-Sec61, *α* (control), HaloTag-KDEL, and 3×HaloTag-KDEL. **E** Population statistics of the effective diffusion coefficient *D*_eff_ for the same conditions.

### D. Dynamics of misfolded proteins in the ER lumen

One of the major functions of the ER is to serve as the major site of translation, folding, and quality control for membrane and secreted proteins [60, 62]. Consequently, much of the ER lumen is dominated by protein-folding machinery, whose interactions with ER-localized cargo are likely to have significant effects on the diffusive properties of the protein species within. Studies using FRAP and FLIP have established dramatic changes in the diffusive properties of ER-resident protein folding machinery under conditions of disregulated folding [36, 37, 69, 72], but they have generally been less effective in resolving changes in the diffusive properties of the misfolded proteins themselves [50], despite evidence from FCS that such changes should be present in the same systems [44] (see *Supplementary Text and Discussion*, Section 4). Additionally, literature examining the diffusive properties of number of unrelated targets has yielded a confusing combination of gains and losses of motility in the unfolded state [49, 59, 61, 64], suggesting a need for a standardized, full-cell approach to characterizing the redistribution of misfolded proteins over time.

To test if the increased resolving power of our approach could address this in principle, we took advantage of the fact that the protein CD3*δ* is obligately misfolded and degraded in normal tissue culture cells when the other components of the CD3 complex are not present [7, 9]. Truncation experiments have shown that the luminal domain alone is sufficient for misfolding and degradation when expressed as a soluble construct [7, 46], so we tested its effect on the motility of freely diffusing HaloTag in the ER lumen. Briey, we fused the signal sequence and soluble domain of CD3*δ* to HaloTag to generate a photoactivatable version of a model misfolded protein (CD3*δ*Δ–HaloTag).The resulting protein is a good client for ER-associated degradation through the Hrd1 pathway, indicating it is correctly recognized as misfolded by the ER quality control system and eventually removed from the ER. As such, we reasoned that it should at the very least switch between diffusive motion in the lumen and motion associated with Hrd1, which is a membrane-embedded protein. We then performed our full analysis pipeline, and analysed the dispersion of CD3*δ*Δ–HaloTag from the PA source over time.

Examination of cells expressing the misfolded construct showed that a significant fraction of the protein was still dynamic within the system, allowing us to perform the full analysis pipeline (fig 5A-B). Notably, CD3*δ*Δ–HaloTag shows significant delayed time of arrival for photoactivated protein compared to a HaloTag-KDEL control (fig 5C), suggesting this approach can resolve slowed diffusive properties of proteins as a result of interactions with luminal protein folding machinery. However, like the other constructs, the majority of the data was well described by a simple diffusion model once the ER structure was accounted for (fig 5D), suggesting the sources of anomality observed in FCS are either at spatiotemporal scales beneath the resolution of this technique or only exist within subpopulations of molecules that do not diffuse across the micron scales visible in these experiments. However, the mean effective diffusion coefficient is significantly reduced for the misfolded protein compared to the HaloTag-KDEL control (*D*_CD3 *δ* Δ_ = 6.19 μm^2^ s^−1^, *D*_KDEL_ = 16.84 μm^2^ s^−1^, p-value < 1 × 10^−3^, see fig 5E), suggesting interactions with ER quality control machinery are frequent enough to create a reduced effective diffusion coefficient across the population (see *Supplementary Text and Discussion*, Section 5). We note that this reduced effective diffusion coefficient is closer to that for membrane-anchored proteins than it is to other luminal content, even though CD3*δ*Δ-HaloTag is signifcantly smaller than the 3xHaloTag-KDEL construct introduced previously (see fig 4), suggesting our approach can identify the known significant interactions with misfolding machinery, even if they occur over spatiotemporal scales too small to resolve them directly.

**Figure 5.**
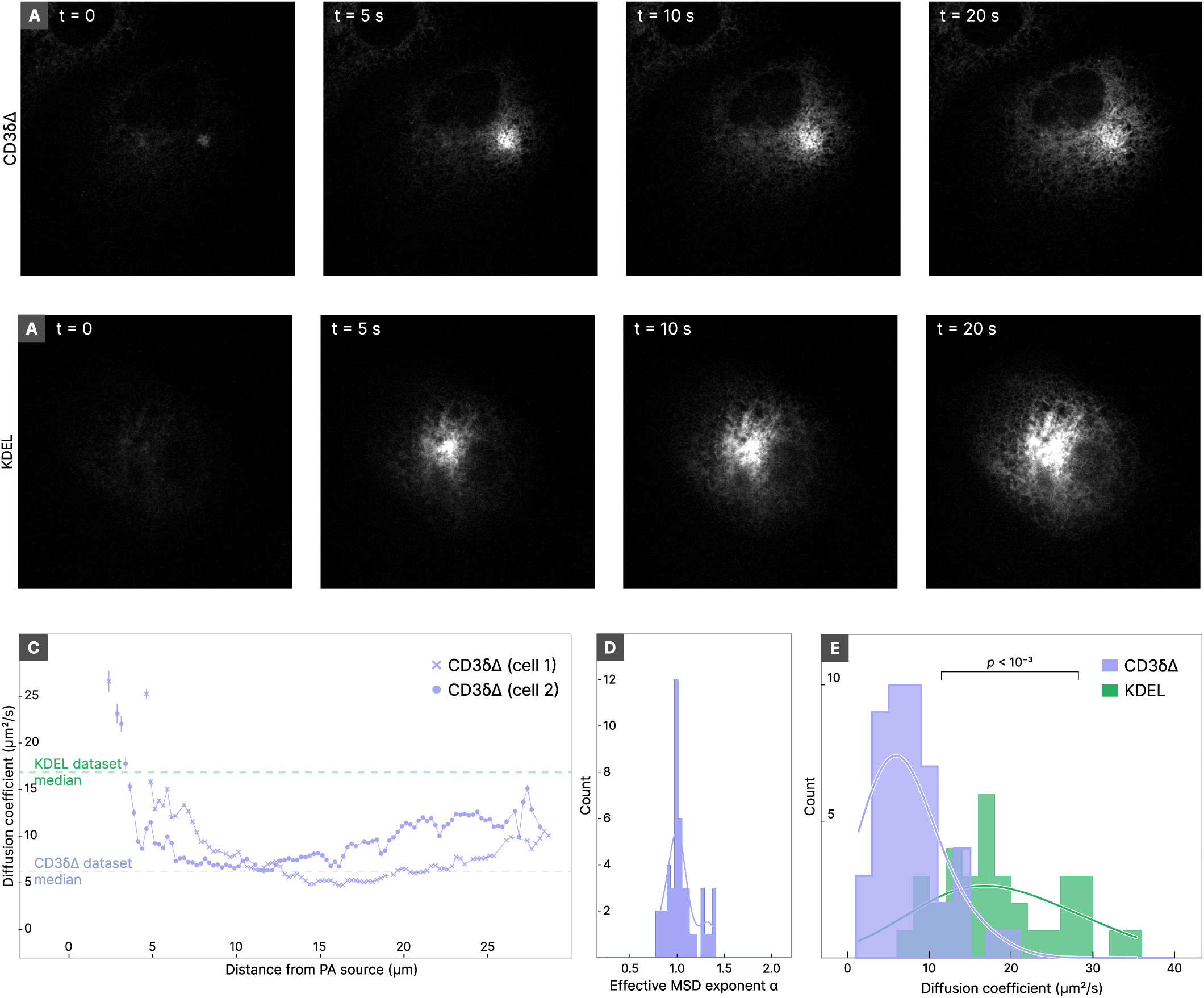
Misfolded luminal proteins show slowed effective diffusion. **A-B** Representative examples of the spatial distribution of photoactivated, misfolded protein (CD3-*δ* Δ-HaloTag)(A) or freely diffusing folded protein (HaloTag-KDEL) (B). **C** Isotropic diffusion fit (averaged over all points at a fixed distance) for two example cells expressing CD3-*δ* Δ-HaloTag overlaid against the median of the population. The median value for cells expressing HaloTag-KDEL is shown for comparison. **D** Distribution of the effective MSD exponent for the cells expressing CD3-*δ* Δ-HaloTag. **E** Distribution of effective diffusion coefficient *D*_eff_ for CD3-*δ* Δ-HaloTag (lavender).The distribution of mean *D*_eff_ for HaloTag-KDEL is reproduced from Figure 1 for comparison.

### E. Spatial heterogeneity in molecular mixing in the ER

One limitation of existing technologies for understanding the dynamics of ER content is their difficulty in identifying spatially varying mixing frequencies or diffusion speeds.The data presented thus far is analysed assuming a reasonable degree of isotropy in the system, since locations at similar distances from the PA source are averaged together. While this approach generally fits the data well, we wondered if the high resolution afforded by the approach might allow us to resolve heterogeneity across the structure or locally varying mixing frequencies.

To address this, we divided each cell into spatially uniform regions (in our case, a square grid) and local regions of the ER within each grid square were binned together and collectively fit for a diffusion model with the PA source signal imposed as a boundary condition (fig 6A). In an unmodified form, this local approach suers from low signal to noise due to background and moving ER structure [71], but our high-speed, two-channel approach and machine learning-assisted structural estimation provided a way to surmount this problem. Within each square on the grid, only pixels within the segmented structure from the moxGFP signal were analyzed (fig 6B), and the pixels used for analysis were updated in real time as the structure moved. Bins that did not contain ER structure for at least half of the frames in the timelapse were discarded. The resulting time evolution curve for each non-empty grid bin was fit as described for averaged distances (e.g., fig 6C), and by fixing the PA source parameters as described above, we estimated an effective diffusion coefficient for each grid bin (see *Methods*, section IV C).The resulting spatially-defined effective diffusion coefficients provided a map of regions in the cellular landscape where diffusion is faster or slower than predicted by the median diffusion coefficient in the cell (fig 6D). Regions characterized by higher effective diffusion coffiefficients are associated with faster arrival of luminal proteins from the PA source.

**Figure 6.**
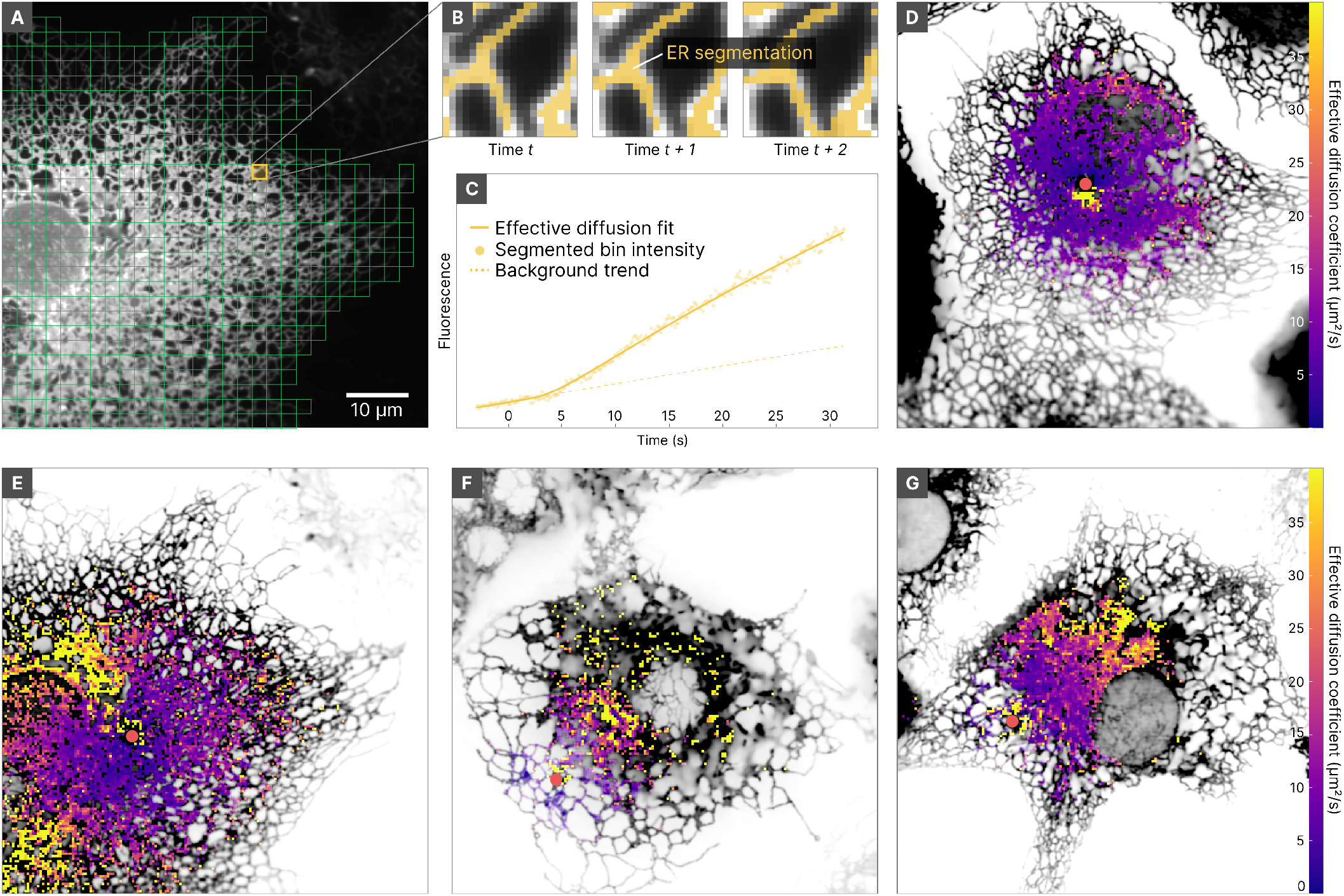
Apparent heterogeneity in luminal mixing dynamics. **A** Subdivision of the ER using a grid of square bins. (Note: a larger bin size is used for easier visualization in the figure than was used for analysis.) **B** An example of ER-associated, segmented pixels belonging to a given bin as they are tracked through time. **C** Average fluorescence over time for segmented pixels belonging to the bin shown in B. Dotted line shows the model fit. **D** Example of a spatial map describing the observed mean arrival time of HaloTag-KDEL in a representative cell, visualized by a spatially-dependent effective diffusion coefficient (higher is faster).The red dot indicates the PA source. The grid bin size is 400 nm (3×3 camera pixels). Note the relatively uniform diffusion throughout the structure. **E-H** Examples of diffusion maps showing asymmetric arrival times in subregions of the ER network. Diffusion maps are derived from cells expressing HaloTag-KDEL (E), 3xHaloTag-KDEL (F), and CD3*δ*Δ-HaloTag (G).

In contrast to the majority of cells which were largely uniform (e.g., fig 6D), in a few of the cells in each condition we observed cases of strong local heterogeneity in the dispersal of fluorescent proteins, represented by regions exhibiting faster arrival time with respect to the cell median mixing time (e.g., fig 6E-G, yellow regions). Such regions seem to most frequently be localized in denser perinuclear regions of the ER, close to the nuclear envelope. To ascertain whether this heterogeneity could be explained by an increased reshaping of the ER morphology in the affected regions, we quantified the morphology variation and tubule motion (see *Methods*, section IV A and supplementary figure S2). ere was no association between higher morphological reshaping and regions with increased mixing dynamics, suggesting that motion of the ER structure is unlikely to be the source of the observed anisotropy in the redistribution timescales.

Although relatively rare, we found examples of such anisotropic photoactivated protein arrival in all of luminal constructs (fig 6E-G), but not the membrane-anchored constructs. The location and size of these regions of faster and slower protein arrival are not consistent with the predicted spatial scales for active transport [16, 31], which are thought to occur over much smaller scales. However, we note that the regions of faster arrival time are in regions of the cell where we have previously shown an elevated level of ER connectivity [84], so they may represent bias in local mixing as a result of the connectivity of the ER network, as has been demonstrated with correlated FRAP and single molecule tracking in systems with more predictable cross sectional geometries [4]. Such a system could potentially give rise the frequently observed but poorly understood phenomena of locally regulated ER functions (see *Supplementary Text and Discussion*, Section 6).

## III. DISCUSSION

In this article, we have combined high speed imaging with point-based photoactivation to examine the dynamic properties of proteins within the ER. We have implemented this approach with a machine learning-based segmentation pipeline and a spatially-defined modeling procedure that can extract quantitative information about molecular motion within the network. This approach supports evaluation of the primary mode of molecular motion at high speed across entire cells, and using this tool we have made a variety of observations about the physical factors that govern molecular behaviour within the ER. Most notably, we find that at the micron-scale, transport phenomena within the ER lumen and membrane are well described by a simple diffusion model when a first degree approximation of the ER structure is accounted for, without needing more complex active ow or anomalous models for diffusion. This model dictates that diffusion within both the membrane and the lumen is likely significantly faster than previous models that do not account for the ER structure, but it does produce effective diffusion coefficients consistent with other structure-based analyses [67, 73] or experiments performed over spatial scales where ER structural contributions are thought to be negligible [49, 78] (see *Supplementary Text and Discussion*, Section 1).

While this result is apparently at odds with models of locally directed flows [16, 31] or anomalous diffusion as a result of local binding [44], we caution that this approach only observes processes visible at the spatial and temporal scales of diffraction-limited imaging over hundreds of milliseconds. Local, nonlinear transport phenomena at the nanoscale can still show linear MSD scaling over larger spatial and temporal scales if the phenomena are locally isotropic [16, 44], a phenomenon we observe directly in the predominantly diffusive motion of misfolded proteins that are known to have local binding interactions with folding machinery (see *Supplementary Text and Discussion*, Section 5). us, we conclude that if local active flows are present, they must exist over spatial and temporal scales that average out to net diffusive behavior by the micron scale, as has been shown for sources of anomality in ER protein diffusion with other models [44, 49].

We also note from this a necessary caution about models in complex structures like the ER. It has been well established by theory [66, 68] and experiment [15, 67, 73] that confinement of diffusing particles to networks can result in the appearance of subdiffusive motion. In this work, which uses full cells that are significantly less uniform in their connectivity than the examples in the literature, we see examples where confinement to the structure has more dramatic effects at short distances and time scales than at larger ones, resulting in the appearance of superdiffusive motion (see *Supplementary Text and Discussion*, Sections 1 and 6 for full discussion).Thus, depending on complexity and connectivity of the underlying structure, MSD scaling can be misleading in either direction about the true nature of the motion within.

The combinatorial approach introduced here promises to be broadly implementable for dealing with complex structured scaffolds on which transport phenomena occur even beyond the ER, and will undoubtedly be improved by integration with developing technologies that can further resolve the complex structures that exist in cells [29, 54, 79]. In particular, one area interest for further technology development is the integration of this approach with tools that have complimentary spatial and temporal scales. For example, traditionally the regionspecific nature of FRAP has made it difficult to multiplex in the same cells or regions as FCS or single molecule tracking, which can in principle be performed simultaneously with our approach. This type of integration across scales at the population level has been effective even applied at the cell population level [4, 44, 49], and promises to elucidate many of the open questions about folding and interaction dynamics within organelles when calibrated inside individual cells.

## IV. METHODS

Fluorescence microscopy images are acquired in low-light conditions, since the power of the excitation laser and exposure interval are constrained by the fluorescence saturation rate, the risk of photodamaging the sample, and the acquisition rate for dynamical imaging. Fluorescent images are thus characterized by high level of noise. In particular, the main source of disturbance is represented by Poisson noise (or *shot noise*) originating from the discrete number of photons hitting the sensor [21]. This is combined with other signal-independent noise sources, due to thermal vibration (e.g. dark current) or caused by the electronics (i.e. noise due to the amplification, conversion, and transmission process, which is usually collectively indicated as *readout noise*).These last noise sources do not depend on the photon flux, and can be modelled effectively as a single aggregated white noise source. Thus, denoising of fluorescence images requires handling Poisson-Gaussian noise. Various techniques have been developed to solve this problem, following mainly three approaches: variance stabilizing transformation followed by additive white noise removal [21], algorithms specifically designed for Poisson-Gaussian mixtures such as PURE-LET [43], and deep learning models based on CNN [38, 80]. A recent work, benchmarking the most popular methods on a dataset of fluorescence images, highlighted the superior performances of CNN-based models [82].The main drawback of CNN is that they usually require training on a set comprising both noisy and clean images. However, in the last few years various deep learning models have been proposed which allow training without clean data [5, 38, 76]. Another drawback of CNNs is the phenomenon of pareidolia, i.e. the tendency to impose a meaningful interpretation to random or ambiguous patterns. CNN are, by their nature, biased towards certain structures which can be intrinsic or derived from the training data [47, 76]. While this is the very reason of their effectiveness, it can also be the cause of unwanted visual artefacts: a CNN model trained on ER images will develop a tendency to see ER even where there is none. For these reasons, in our analysis of photoactivation data, we have taken a mixed approach: we relied on a deep-learning model to facilitate the segmentation of the ER morphology, but followed a more conservative approach when extracting quantitative measures from the photoactivation curves, considering a statistical model of the raw data.

### A. Pre-processing and segmentation of ER morphology

To ensure reliable segmentation of the ER structure, we preprocess the data of the morphology channel (GFP channel) in several steps (see supplementary fig S1).Thse preprocessing steps are needed because the ER structure and density can vary significantly. Regions of the ER close to the nuclear envelope are characterized by denser structures (membrane sheets or dense matrices of tubules [53, 74]) which are not resolvable with confocal microscopy and thus appear as continuous membrane arrangements. At the cell periphery, the ER is formed by a network of sparse tubules with diameter 60–100 nm [53, 70].These structural differences determine the quantity of fluorescent proteins and worsen the image contrast, making it challenging to segment simultaneously the entire ER structure [56].

#### 1. Frame denoising

We first denoised the morphology data to reduce the level of noise and improve the detection of finer structures. To this aim, we adapted the Noise2Noise technique [38] to train a U-Net convolutional neural network [63] for noise removal of single frames (see supplementary fig. S1A). We trained U-Net model using pairs of noisy realizations generated by extracting 128×128 px tiles from time-adjacent frames. One noisy realization is kept as a reference while the other is processed by the U-Net model. The loss is then calculated as the MSE between the reference noisy frame and the model output. This approach relies on the assumption that the acquisition rate is fast enough to make changes between consecutive frames negligible.The assumption is reasonable as most of the ER structure is steady for the acquisition interval used (100 ms). To minimize the bias created by the time evolution of the ER network, both the order of the training samples and the order of the two images forming a couple were randomized in the repeated training epochs. We kept 10% of the samples for validation, separating them from training data.The training was performed on batches of 4 images, using the Adam optimization algorithm [34] with parameters *β*_1_ = 0.9, *β*_2_ = 0.99, no weight decay and initial learning rate 1 = 2.75 10^*-*4^. The learning rate was then gradually reduced with a factor 0.75 when the MSE loss calculated on the validation set did not improve over 5 epochs. To prevent overing, training was stopped when the loss over the validation set did not improve for 15 consecutive epochs, selecting the model with minimal loss.

#### 2. Sharpening and structure enhancement

To enhance the resolution of the finer ER structure, we deconvolved the morphology frames to create sharper images. Image deconvolution is an inverse problem of the type ***y*** = **A*x***, consisting in recovering the sharp image ***x*** from a blurred version ***y***, with **A** representing the PSF operator associated to the optical system. Since the problem is ill-posed, the solution is found by adding regularization constraints on ***x*** such as total variation [23]. Recently, a sparsity constraint in the deconvolution procedure has been shown to successfully increase the spatio-temporal resolution in fluorescence microscopy [83]. Based on these approaches, we defined a spatio-temporal deconvolution problem with total variation and sparsity regularization terms, to provide good reconstruction of edges, and a smoothness constraint in time dimension to account the gradual structural movements. We define the optimization problem as

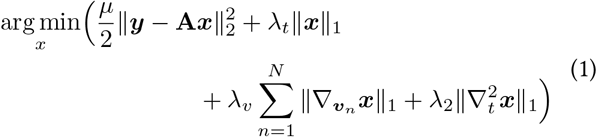

where y represents a sequence of frames, A is the PSF operator, ‖*·* ‖ and 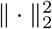 represent respectively the ℓ_1_ and ℓ_2_ norms, 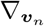 is the directional derivative along ***v***_*n*_, and 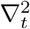 is the second derivative in the time dimension. Multiple spatial directions ***v***_*n*_ = (sin(2π*n*/*N*), cos(2π*n*/*N*), 0) in space (*x, y, t*) to reduce staircase artefacts. To ensure correct enforcement of the sparsity constraint, each frame in ***y*** was preprocessed by taking a 2D discrete wavelet transform and dampening the approximation coefficients by a factor 0.5. The operator A was assumed to be a Gaussian PSF with appropriate standard deviation (1.27 px).The solution ***x*** was computed with the split Bregman method [24]. Parameter values used in the optimization are reported in supplementary table S1.

#### 3. Segmentation of the network structure

The ER network was segmented from the sharpened stack as follows. First, the frames were clipped at zero and rescaled between 0 and 1 and local contrast was enhanced by contrast limited adaptive histogram equalization (CLAHE) [1] for five iterations. To detect tubular structures, the frames were first transformed with a Sato tubeness filter [65] with σ = 100 nm and then binarized by locally adaptive Niblack thresholding [51] with window size 127 px (≈ 17 m) and weight 0.1. Empty background regions, which could create bad detections with purely local thresholding, were excluded by setting a minimum global threshold. Continuous surfaces corresponding to dense matrices and cisternae were obtained by a similar thresholding procedure, followed by a morphological opening operation with a disk of radius 250 nm. The thresholded tubules and sheets were then merged in a single frame sequence. To preserve details in very close tubular structure, local minima were identified with a h-minimum transform (with threshold 1 × 10^−3^) before and set to zero [56]. To fix occasional flickering of isolated pixels, a morphological closing operation was applied along the time axis of the thresholded stack. Finally, the network structure was extracted by thinning procedure [81]. To preserve dense matrices and cisternæ, these regions were again extracted by morphological opening and added back to the skeletonized network structure.

### B. Estimation of morphology reshaping

The amount of local morphological reshaping was quantified in two ways. First, by estimating the relative variance of the denoised morphology frames along the time stack, highlighting regions of the network undergoing more variation in time. Predictably, most of the variation was localized on the tubule edges, due to small oscillations (fig S2B, blue shade). Note that in our method, such small oscillations do not affect the estimate of the dynamics, as only the pixels clearly belonging to the ER structure are considered. Second, we estimated motion of tubules and other structures by optical ow using Farnebäck’s algorithm [19], then calculated the mean norm of the velocity vectors over the photoactivation time to reveal regions characterized by more active reshaping of the network structure (fig S2B, red shade).

### C. Statistical model of the photoactivatable channel

#### 1. Estimation of the noise model parameters

We considered a Poisson-Gaussian noise model with clipping as described in [20, 21].The original unclipped noise model which describes the observed pixel value z (x) at position x is defined by

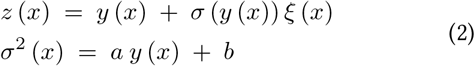

where y (x) is the true pixel value, *ξ* (x) is zero-mean and unit variance random noise, and σ (y) is the y-dependent standard deviation function characterizing the Poisson-Gaussian noise. Note that this model assumes that the photon count y is large enough, such that the Poisson term can be approximated with a normally distributed noise with signal-dependent variance. Imaging sensors are also characterized by a bias level *µ*, i.e. an artificial offset added to the collected charge to reduce clipping. This offset value must be subtracted from the readout to correctly model the signal-dependent part of the noise, e.g. by considering the variance function σ (*y* − µ) or by redefining y←y−µ. If the sensor is not able to capture the full dynamic range of the image (particularly in the case of fluorescence imaging, where frames are intrinsically underexposed), one can define a clipped observation model as:

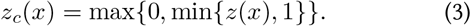

The goal is to reconstruct the model parameters a, b, µ from the video data. We applied the technique described in [21] on all frames, segmenting the full video data in level sets, excluding the tubules boundaries obtained from the segmentation of the morphology channel, to estimate 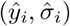 pairs. We then performed a maximum likelihood t of the model parameters using the L-BFGS algorithm [48, 86].

#### 2. Effective diffusion model

To extract quantitative features describing the protein dynamics, we consider an effective 2D diffusion model. The approximation with a 2D space is appropriate given that the ER structure in COS-7 cells can, for most of its surface, described by a planar graph [53]. While it is possible to derive a reaction-diffusion system which modelling a constant-rate photoconversion in the source ROI, this approach requires knowledge (or t) of the photo conversion rate parameter. Moreover, due to the reshaping of the ER, the ROI exposed to the photoactivation laser may change constantly, altering the effectiveness of photo conversion. To avoid these problems, we used a hybrid approach which allows to preemptively tune the diffusion model based on the data. To this aim, we considered a diffusion model in which we impose a time-dependent Dirichlet boundary condition corresponding to the fluorescence measured close to the photoactivation ROI. We denote by *ϕ*(*t*) the protein concentration measured at distance *R* from the centre of the photoactivation ROI. Given an analytical expression for ϕ(*t*), we can calculate the solution of the radial diffusion equation *u*(*r, t*) (see supplementary IV A, eq. S8):

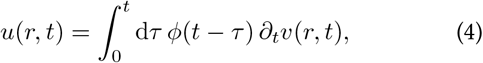

where *v*(*r, t*) is the solution of the auxiliary diffusion problem with a constant boundary condition (*ϕ*(*t*)= 1), which can be explicitly expressed in Laplace domain as

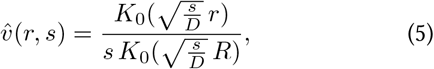

where *K*_0_ is the modified Bessel function of order zero. It is convenient to express the convolution in eq. (4) as a product in the Laplace domain:

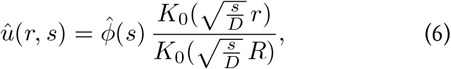

calculating the *u*(*r, t*) by inverting the Laplace transformation.

In principle, we can choose any analytical form of *ϕ*(*t*) that well describes the fluorescence curve at distance *R* from the centre of the photoactivation ROI observed in the experimental data. We considered here the particular case where *ϕ* (*t*) can be expressed as a sum of exponentially saturating functions:

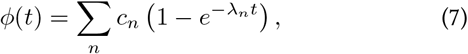

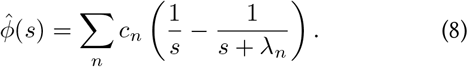

Using this definition for *ϕ* combined with eq. 6 we can write

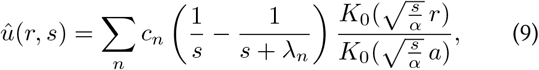

from which *u*(*r, t*) can be obtained by inverting the Laplace transform numerically.

#### 3. Fluorescence intensity model

To properly describe the fluorescence data, we need to take into account variations which not related to the diffusion process. These are mainly constituted by the background intensity and the random photoconversion of proteins outside the photoactivation region due to leaks from the other channel and acquisition laser. We describe these by an offset and a linear trend which can be observed before the photoactivation laser is activated.Thus, we describe the fluorescence intensity *f*(**x**, *t*) at position ***x*** as the sum of a linear polynomial c_1_ t+c_0_ and the diffusion process *u*(*r, t*):

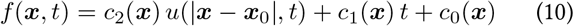

where *x*_0_ is the location of the PA ROI, *c*_0_(***x***) describes the background luminosity, *c*_1_(***x***) is the slope of the linear trend of random photoactivation, and *c*_2_(***x***) is a proportionality constant representing the density of photoactivated proteins. Note that, in general, the coefficients *c*_0_, *c*_1_, and *c*_2_ are spacedependent.

#### 4. Maximum-likelihood t of the diffusion model

We first performed image registration on the photoactivation channel to match the position of the GFP channel by cross-correlation [3, 27]. We then considered the intensity data obtained from the raw images in the form of points *x*_*i*_ = (*r*_*i*_, *z*_*i*_, *t*_*i*_), where r_*i*_ represents the distance from the photoactivation source and z_*i*_ its measured fluorescence at time *t*_*i*_ from the beginning of photoactivation. We used the noise model defined in eq. (2), with parameters determined according to the procedure described above, to estimate the likelihood of the fluorescence model of eq. (10) depending on the *x*_*i*_ samples. e noise variance for a point *x*_*i*_ = (*r*_*i*_, *z*_*i*_, *t*_*i*_) is given by

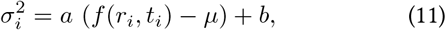

where *f*(*r*_*i*_, *t*_*i*_) is the true fluorescence value and µ is the imaging sensor bias level. We consider 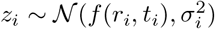 The log-likelihood is then

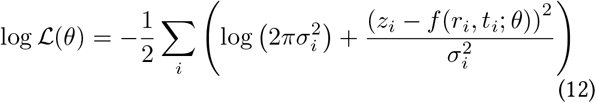

with θ is the set of parameters *c*_0_, *c*_1_, *c*_2_, *D* (see eq. (10)).The maximum likelihood parameter is obtained as

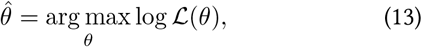

which can be solved by numerical optimization. This maximum likelihood estimation can be extended to ensemble points obtained by averaging multiple pixel values in the photoactivation stack. The noise variance of such aggregated points is simply

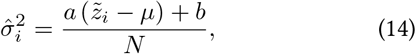

where *N* is the number of averaged pixels and 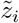 is average fluorescence value. By maximum-likelihood estimation it is then possible to recover the diffusion coefficient *D* and the coefficients 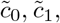, and 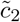 corresponding respectively to the average background intensity, average slope of the linear trend, and average density over the ensemble. In practice, we t the model parameters using L-BFGS-B [48, 86] in two separate steps. First we performed the maximum likelihood fit on the interval preceding photoactivation, recovering the trend para-meters *c*_1_ and *c*_0_.Then, fixing *c*_1_ and *c*_0_, we performed the fit on the full time interval to recover *D* and *c*_2_. We initialized the parameter *c*_2_ based on the GFP-labeled morphology channel as follows. We considered the frames of the morphology channel preceding photoactivation and calculate the average intensities ρ_s_, ρ_t_ and variances 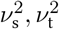 over the pixels corresponding to the PA source and the target regions respectively. We then defined the initial value for coefficient c_2_ as:

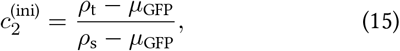

where *µ*_GFP_ is the sensor bias level for the GFP channel. In the morphology-aware approach, we constrained the coefcient *c*_2_ in the interval 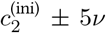, where *v*^2^ is the approximated variance of the ratio distribution obtained as 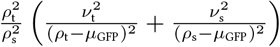.

In the case of isotropic diffusion estimation, these ensemble points were built by binning pixels in each frame based on their distance from the PA source (with bin size of 250 nm). Only pixels belonging to the segmented ER structure were considered. Ensemble points characterized by the same distance *r* were grouped to build a time series 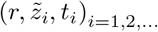. We note that, due to changes in the morphology of the ER, the pixels belonging to a given distance bin are not invariant across frames. This frame by frame adjustment allows cancelling at least part of the effect of the morphology reshaping.

For the estimation of space-dependent effective diffusion, ensemble points were built by dividing the field of view in equal bins using a squared grid (fig 6A). For each bin, average fluorescence intensity and median distance from the source were evaluated in each frame (considering only the pixels belonging to the segmented ER structure, fig 6B). The ensemble points obtained by this procedure were then used to estimate the effective diffusion by maximum likelihood (fig 6C), obtaining a diffusion coefficient for each bin of the grid (fig 6D).

We estimated the variance of the fitted parameter *D* using the observed Fisher information. In the limit of large number of samples, the distribution of the maximum likelihood estimate 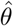 can be approximated as [28]:

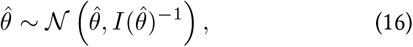

where 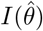 is the observed Fisher information. In our case, the observed Fisher information can be evaluated as the Hessian of the negative log-likelihood defined in eq. (12), i.e.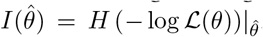. We thus estimated the variance of the parameter D by numerically evaluating the inverse Hessian of log at its minimum [10], and taking the appropriate diagonal element. We note that such estimate of the error is statistically meaningful only limited to the context of the optimization problem defined in eq. (13), as it does not take into account errors resulting from inappropriateness of the effective diffusion model or bad ER segmentation. Nevertheless, this approach provides a method to systematically rate the quality of the ts and exclude the unreliable datapoints in the estimation of the effective diffusion coefficient (fig 1C).

## References

[1] The Graphics Gems Series. In Paul S. Heckbert, editor, Graphics Gems, page ii. Academic Press.

[2] Bastian R Angermann, Frederick Klauschen, Alex D Garcia, Thorsten Prustel, Fengkai Zhang, Ronald N Germain, and Martin Meier-Schellersheim. Computational modeling of cellular signaling processes embedded into dynamic spatial contexts. Nature methods, 9(3):283–289, 2012.

[3] Paul E Anuta. Spatial registration of multispectral and multitemporal digital imagery using fast fourier transform techniques. IEEE transactions on Geoscience Electronics, 8(4):353–368, 1970.

[4] Somenath Bakshi, Benjamin P Bratton, and James C Weisshaar. Subdiffraction-limit study of kaede diffusion and spatial distribution in live escherichia coli. Biophysical journal, 101(10):2535–2544, 2011.

[5] Joshua Batson and Loic Royer. Noise2Self: Blind denoising by self-supervision. In International Conference on Machine Learning, pages 524–533.

[6] Buzz Baum and David A Baum. The merger that made us. BMC biology, 18(1):1–4, 2020.

[7] Riccardo Bernasconi, Carmela Galli, Verena Calanca, Toshihiro Nakajima, and Maurizio Molinari. Stringent requirement for hrd1, sel1l, and os-9/xtp3-b for disposal of erad-ls substrates. Journal of Cell Biology, 188(2):223–235, 2010.

[8] Michael J Berridge. The endoplasmic reticulum: a multifunctional signaling organelle. Cell calcium, 32(5-6):235–249, 2002.

[9] Juan S Bonifacino, Carolyn K Suzuki, Jennifer Lippincotttt-Schwartz, Allan M Weissman, and Richard D Klausner. Pregolgi degradation of newly synthesized t-cell antigen receptor chains: intrinsic sensitivity and the role of subunit assembly. The Journal of cell biology, 109(1):73–83, 1989.

[10] Per A. Brodtkorb and John D’Errico. numdiools. https://github.com/pbrod/numdifftools, 2015.

[11] Nelson B Cole, Carolyn L Smith, Noah Sciaky, Mark Terasaki, Michael Edidin, and Jennifer Lippincotttt-Schwartz. Diffusional mobility of golgi proteins in membranes of living cells. Science, 273(5276):797–801, 1996.

[12] Lindsey M Costantini, Mikhail Baloban, Michele L Markwardt, Megan A Rizzo, Feng Guo, Vladislav V Verkhusha, and Erik L Snapp. A palette of fluorescent proteins optimized for diverse cellular environments. Nature communications, 6(1):1–13, 2015.

[13] Lindsey M Costantini and Erik Lee Snapp. Fluorescent proteins in cellular organelles: serious pitfalls and some solutions. DNA and cell biology, 32(11):622–627, 2013.

[14] Sándor Damjanovich. Mobility and proximity in biological membranes. CRC Press, 2018.

[15] Mark J Dayel, Erik FY Hom, and Alan S Verkman. Diffusion of green fluorescent protein in the aqueous-phase lumen of endoplasmic reticulum. Biophysical journal, 76(5):2843–2851, 1999.

[16] M Dora and D Holcman. Active flow network generates molecular transport by packets: case of the endoplasmic reticulum. Proceedings of the Royal Society B, 287(1930):20200493, 2020.

[17] Jan Ellenberg, Jennifer Lippincotttt-Schwartz, and John F Presley. Dual-colour imaging with gfp variants. Trends in cell biology, 9(2):52–56, 1999.

[18] Lance P Encell, Rachel Friedman Ohana, Kris Zimmerman, Paul Otto, Gediminas Vidugiris, Monika G Wood, Georgyi V Los, Mark G McDougall, Chad Zimprich, Natasha Karassina, et al. Suppl 1: Development of a dehalogenase-based protein fusion tag capable of rapid, selective and covalent attachment to customizable ligands. Current chemical genomics, 6:55, 2012.

[19] Gunnar Farnebäck. Two-frame motion estimation based on polynomial expansion. In Scandinavian conference on Image analysis, pages 363–370. Springer, 2003.

[20] Alessandro Foi. Clipped noisy images: Heteroskedastic modeling and practical denoising. 89(12):2609–2629.

[21] Alessandro Foi, Mejdi Trimeche, Vladimir Katkovnik, and Karen Egiazarian. Practical Poissonian-Gaussian noise modeling and fitting for single-image raw-data. 17(10):1737–1754.

[22] Larry D Frye and Michael Edidin. The rapid intermixing of cell surface antigens after formation of mouse-human heterokaryons. Journal of cell science, 7(2):319–335, 1970.

[23] Pascal Getreuer. Total Variation Deconvolution using Split Bregman. 2:158–174.

[24] Tom Goldstein and Stanley Osher. The Split Bregman Method for L1-Regularized Problems. 2(2):323–343.

[25] Uma Goyal and Craig Blackstone. Untangling the web: mechanisms underlying er network formation. Biochimica et Biophysica Acta (BBA)-Molecular Cell Research, 1833(11):2492–2498, 2013.

[26] Jonathan B Grimm, Brian P English, Heejun Choi, Anand K Muthusamy, Brian P Mehl, Peng Dong, Timothy A Brown, Jennifer Lippincott-Schwartz, Zhe Liu, Timothée Lionnet, et al. Bright photoactivatable uorophores for single-molecule imaging. Nature methods, 13(12):985–988, 2016.

[27] Manuel Guizar-Sicairos, Samuel Turman, and James R Fienup. Efficient subpixel image registration algorithms. Optics letters, 33(2):156–158, 2008.

[28] Trevor Hastie, Robert Tibshirani, Jerome H Friedman, and Jerome H Friedman. The elements of statistical learning: data mining, inference, and prediction, volume 2. Springer, 2009.

[29] Larissa Heinrich, Davis Bennett, David Ackerman, Woohyun Park, John Bogovic, Nils Eckstein, Alyson Petruncio, Jody Clements, Song Pang, C Shan Xu, et al. Whole-cell organelle segmentation in volume electron microscopy. Nature, 599(7883):141–146, 2021.

[30] Koret Hirschberg, Chad M Miller, Jan Ellenberg, John F Presley, Eric D Siggia, Robert D Phair, and Jennifer Lippincotttt-Schwartz. Kinetic analysis of secretory protein traffic and characterization of golgi to plasma membrane transport intermediates in living cells. The Journal of cell biology, 143(6):1485–1503, 1998.

[31] David Holcman, Pierre Parutto, Joseph E. Chambers, Marcus Fantham, Laurence J. Young, Stefan J. Marciniak, Clemens F. Kaminski, David Ron, and Edward Avezov. Single particle trajectories reveal active endoplasmic reticulum luminal ow. 20(10):1118–1125.

[32] Maximilian AH Jakobs, Andrea Dimitracopoulos, and Kristian Franze. Kymobutler, a deep learning software for automated kymograph analysis. Elife, 8:e42288, 2019.

[33] Andrii A Kaberniuk, Manuel A Mohr, Vladislav V Verkhusha, and Erik Lee Snapp. moxmaple3: a photoswitchable fluorescent protein for palm and protein highlighting in oxidizing cellular environments. Scientific reports, 8(1):1–10, 2018.

[34] Diederik P. Kingma and Jimmy Ba. Adam: A Method for Stochastic Optimization.

[35] Tasuku Konno, Pierre Parutto, David MD Bailey, Valentina Daví, Cécile Crapart, Mosab Ali Awadelkareem, Colin Hockings, Aidan Brown, Katherine M Xiang, Anamika Agrawal, et al. Endoplasmic reticulum morphological regulation by rtn4/nogo modulates neuronal regeneration by curbing luminal transport. bioRxiv, pages 2021–05, 2021.

[36] Chun Wei Lai, Deborah E Aronson, and Erik Lee Snapp. Bip availability distinguishes states of homeostasis and stress in the endoplasmic reticulum of living cells. Molecular biology of the cell, 21(12):1909–1921, 2010.

[37] Chunwei Walter Lai, Joel H Otero, Linda M Hendershot, and Erik Snapp. Erdj4 protein is a soluble endoplasmic reticulum (er) dnaj family protein that interacts with er-associated degradation machinery. Journal of Biological Chemistry, 287(11):7969–7978, 2012.

[38] Jaakko Lehtinen, Jacob Munkberg, Jon Hasselgren, Samuli Laine, Tero Karras, Miika Aittala, and Timo Aila. Noise2Noise: Learning image restoration without clean data. volume 80 of Proceedings of Machine Learning Research, pages 2965–2974. PMLR.

[39] Jennifer Lippincott-Schwartz, Nihal Altan-Bonnet, and George H Patterson. Photobleaching and photoactivation: Following protein dynamics in living cells. pages S7–14.

[40] Jennifer Lippincott-Schwartz, Erik Snapp, and Anne Kenworthy. Studying protein dynamics in living cells. Nature reviews Molecular cell biology, 2(6):444–456, 2001.

[41] Jennifer Lippincott-Schwartz, Erik Lee Snapp, and Robert D. Phair. e Development and Enhancement of FRAP as a Key Tool for Investigating Protein Dynamics. 115(7):1146–1155.

[42] Georgyi V Los, Lance P Encell, Mark G McDougall, Danette D Hartzell, Natasha Karassina, Chad Zimprich, Monika G Wood, Randy Learish, Rachel Friedman Ohana, Marjeta Urh, et al. Halotag: a novel protein labeling technology for cell imaging and protein analysis. ACS chemical biology, 3(6):373–382, 2008.

[43] Florian Luisier, ierry Blu, and Michael Unser. Image Denoising in Mixed Poisson–Gaussian Noise. 20(3):696–708.

[44] Nina Malchus and Matthias Weiss. Anomalous diffusion reports on the interaction of misfolded proteins with the quality control machinery in the endoplasmic reticulum. Biophysical journal, 99(4):1321–1328, 2010.

[45] Pierre Mangeol, Bram Prevo, and Erwin JG Peterman. Kymographclear and kymographdirect: two tools for the automated quantitative analysis of molecular and cellular dynamics using kymographs. Molecular biology of the cell, 27(12):1948–1957, 2016.

[46] Martin Mehnert, Thomas Sommer, and Ernst Jarosch. Erad ubiquitin ligases: multifunctional tools for protein quality control and waste disposal in the endoplasmic reticulum. Bioessays, 32(10):905–913, 2010.

[47] Benjamin R. Mitchell. e spatial inductive bias of deep learning.

[48] José Luis Morales and Jorge Nocedal. Remark on “algorithm 778: L-bfgs-b: Fortran subroutines for large-scale bound constrained optimization”. ACM Transactions on Mathematical Software (TOMS), 38(1):1–4, 2011.

[49] Hisao Nagaya, Taku Tamura, Arisa Higa-Nishiyama, Koji Ohashi, Mayumi Takeuchi, Hitoshi Hashimoto, Kiyotaka Hatsuzawa, Masataka Kinjo, Tatsuya Okada, and Ikuo Wada. Regulated motion of glycoproteins revealed by direct visualization of a single cargo in the endoplasmic reticulum. The Journal of Cell Biology, 180(1):129–143, 2008.

[50] Sarah Nehls, Erik L Snapp, Nelson B Cole, Kristien JM Zaal, Anne K Kenworthy, Theresa H Roberts, Jan Ellenberg, John F Presley, Eric Siggia, and Jennifer Lippincott-Schwartz. Dynamics and retention of misfolded proteins in native er membranes. Nature cell biology, 2(5):288–295, 2000.

[51] Wayne Niblack. An Introduction to Digital Image Processing. Strandberg Publishing Company.

[52] Benjamin J Nichols, Anne K Kenworthy, Roman S Polishchuk, Robert Lodge, Theresa H Roberts, Koret Hirschberg, Robert D Phair, and Jennifer Lippincott-Schwartz. Rapid cycling of lipid raft markers between the cell surface and golgi complex. The Journal of cell biology, 153(3):529–542, 2001.

[53] Jonathon Nixon-Abell, Christopher J Obara, Aubrey V Weigel, Dong Li, Wesley R Legant, C Shan Xu, H Amalia Pasolli, Kirsten Harvey, Harald F Hess, Eric Betzig, et al. Increased spatiotemporal resolution reveals highly dynamic dense tubular matrices in the peripheral ER. 354(6311):aaf3928.

[54] Christopher J Obara, Andrew S Moore, and Jennifer Lippincott-Schwartz. Structural diversity within the endoplasmic reticulum—from the microscale to the nanoscale. Cold Spring Harbor Perspectives in Biology, page a041259, 2022.

[55] Bence P Ölveczky and AS Verkman. Monte carlo analysis of obstructed diffusion in three dimensions: application to molecular diffusion in organelles. Biophysical journal, 74(5):2722–2730, 1998.

[56] Charloe Pain, Verena Kriechbaumer, Maike Kittelmann, Chris Hawes, and Mark Fricker. Quantitative analysis of plant ER architecture and dynamics. 10(1):984.

[57] George H Patterson and Jennifer Lippincott-Schwartz. A photoactivatable gfp for selective photolabeling of proteins and cells. Science, 297(5588):1873–1877, 2002.

[58] Mu-Ming Poo and Richard A Cone. Lateral diffusion of rhodopsin in the photoreceptor membrane. Nature, 247(5441):438–441, 1974.

[59] Rowena R Ramos, Andrea J Swanson, and Joseph Bass. Calreticulin and hsp90 stabilize the human insulin receptor and promote its mobility in the endoplasmic reticulum. Proceedings of the National Academy of Sciences, 104(25):10470–10475, 2007.

[60] Tom A Rapoport, Long Li, and Eunyong Park. Structural and mechanistic insights into protein translocation. Annual review of cell and developmental biology, 33:369–390, 2017.

[61] Andrew Ridsdale, Maxime Denis, Pierre-Yves Gougeon, Johnny K Ngsee, John F Presley, and Xiaohui Zha. Cholesterol is required for efficient endoplasmic reticulum-to-golgi transport of secretory membrane proteins. Molecular biology of the cell, 17(4):1593–1605, 2006.

[62] David Ron and Peter Walter. Signal integration in the endoplasmic reticulum unfolded protein response. Nature reviews Molecular cell biology, 8(7):519–529, 2007.

[63] Olaf Ronneberger, Philipp Fischer, and omas Brox. U-net: Convolutional networks for biomedical image segmentation. In Lecture Notes in Computer Science, pages 234–241. Springer International Publishing.

[64] Heiko Runz, Kota Miura, Matthias Weiss, and Rainer Pepperkok. Sterols regulate er-export dynamics of secretory cargo protein ts-o45-g. e EMBO journal, 25(13):2953–2965, 2006.

[65] Yoshinobu Sato, Shin Nakajima, Nobuyuki Shiraga, Hideki Atsumi, Shigeyuki Yoshida, omas Koller, Guido Gerig, and Ron Kikinis. ree-dimensional multi-scale line filter for segmentation and visualization of curvilinear structures in medical images. Medical Image Analysis, 2(2):143–168.

[66] Ivo F Sbalzarini, Arnold Hayer, Ari Helenius, and Petros Koumoutsakos. Simulations of (an) isotropic diffusion on curved biological surfaces. Biophysical journal, 90(3):878–885, 2006.

[67] Ivo F Sbalzarini, Anna Mezzacasa, Ari Helenius, and Petros Koumoutsakos. Eects of organelle shape on fluorescence recovery aer photobleaching. Biophysical journal, 89(3):1482–1492, 2005.

[68] Zubenelgenubi C Scott, Aidan I Brown, Saurabh S Mogre, Laura M Westrate, and Elena F Koslover. Diusive search and trajectories on tubular networks: a propagator approach. e European Physical Journal E, 44(6):80, 2021.

[69] Jingshi Shen, Erik L Snapp, Jennifer Lippincott-Schwartz, and Ron Prywes. Stable binding of atf6 to bip in the endoplasmic reticulum stress response. Molecular and cellular biology,25(3):921–932, 2005.

[70] Yoko Shibata, Gia K. Voeltz, and Tom A. Rapoport. Rough sheets and smooth tubules. 126(3):435–439.

[71] Eric D Siggia, Jennifer Lippincott-Schwartz, and Stefan Bekiranov. Diffusion in inhomogeneous media: theory and simulations applied to whole cell photobleach recovery. Biophysical journal, 79(4):1761–1770, 2000.

[72] Erik L Snapp, Ajay Sharma, Jennifer Lippincott-Schwartz, and Ramanujan S Hegde. Monitoring chaperone engagement of substrates in the endoplasmic reticulum of live cells. 103(17):6536–6541.

[73] Yunhao Sun, Zexi Yu, Christopher J. Obara, Keshav Mittal, Jennifer Lippincott-Schwarz, and Elena F. Koslover. Unraveling Single-Particle Trajectories Confined in Tubular Networks.

[74] Mark Terasaki, Lan Bo Chen, and Keigi Fujiwara. Microtubules and the endoplasmic reticulum are highly interdependent structures. 103(4):1557–1568.

[75] Roger Y Tsien. e green uorescent protein. Annual review of biochemistry, 67(1):509–544, 1998.

[76] Dmitry Ulyanov, Andrea Vedaldi, and Victor Lempitsky. Deep Image Prior. pages 9446–9454.

[77] Gerrit Van Meer, Dennis R Voelker, and Gerald W Feigenson. Membrane lipids: where they are and how they behave. Nature reviews Molecular cell biology, 9(2):112–124, 2008.

[78] Mahias Weiss, Hitoshi Hashimoto, and Tommy Nilsson. Anomalous protein diffusion in living cells as seen by fluorescence correlation spectroscopy. Biophysical journal, 84(6):4043–4052, 2003.

[79] C Shan Xu, Song Pang, Gleb Shtengel, Andreas Müller, Alex T Rier, Huxley K Homan, Shin-ya Takemura, Zhiyuan Lu, H Amalia Pasolli, Nirmala Iyer, et al. An open-access volume electron microscopy atlas of whole cells and tissues. Nature, 599(7883):147–151, 2021.

[80] Kai Zhang, Wangmeng Zuo, Yunjin Chen, Deyu Meng, and Lei Zhang. Beyond a Gaussian Denoiser: Residual Learning of Deep CNN for Image Denoising. 26(7):3142–3155.

[81] Tongjie Y. Zhang and Ching Y. Suen. A fast parallel algorithm for thinning digital patterns. 27(3):236–239.

[82] Yide Zhang, Yinhao Zhu, Evan Nichols, Qingfei Wang, Siyuan Zhang, Cody Smith, and Scott Howard. A poisson-gaussian denoising dataset with real fluorescence microscopy images. In CVPR.

[83] Weisong Zhao, Shiqun Zhao, Liuju Li, Xiaoshuai Huang, Shijia Xing, Yulin Zhang, Guohua Qiu, Zhenqian Han, Yingxu Shang, De-en Sun, Chunyan Shan, Runlong Wu, Lusheng Gu, Shuwen Zhang, Riwang Chen, Jian Xiao, Yanquan Mo, Jianyong Wang, Wei Ji, Xing Chen, Baoquan Ding, Yanmei Liu, Heng Mao, Bao-Liang Song, Jiubin Tan, Jian Liu, Haoyu Li, and Liangyi Chen. Sparse deconvolution improves the resolution of live-cell super-resolution fluorescence microscopy. pages 1–12.

[84] Pengli Zheng, Christopher J Obara, Ewa Szczesna, Jonathon Nixon-Abell, Kishore K Mahalingan, Antonina Roll-Mecak, Jennifer Lippincott-Schwartz, and Craig Blackstone. Er proteins decipher the tubulin code to regulate organelle distribution. Nature, 601(7891):132–138, 2022.

[85] H Mimi Zhou, Ingrid Brust-Mascher, and Jonathan M Scholey. Direct visualization of the movement of the monomeric axonal transport motor unc-104 along neuronal processes in livingcaen-orhabditis elegans. Journal of Neuroscience, 21(11):3749–3755, 2001.

[86] Ciyou Zhu, Richard H Byrd, Peihuang Lu, and Jorge Nocedal. Algorithm 778: L-bfgs-b: Fortran subroutines for large-scale bound-constrained optimization. ACM Transactions on mathematical Software (TOMS), 23(4):550–560, 1997.

