## Supplementary Information for "Simultaneous photoactivation and high-speed structural tracking reveal diffusion-dominated motion in the endoplasmic reticulum"

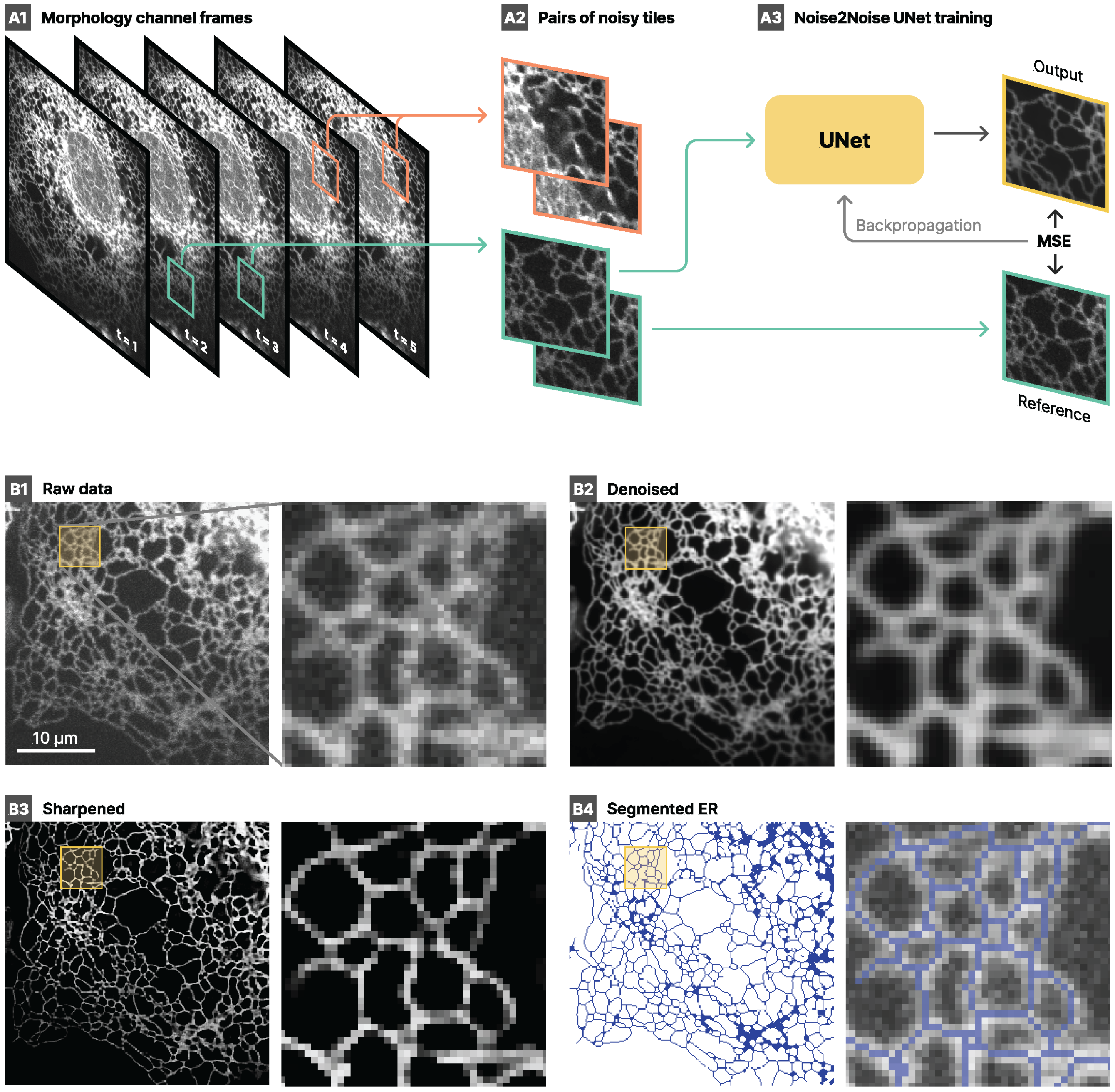


Figure S1. **Morphology processing pipeline.** **A** Data denoising with Noise2Noise method. From the stack of frames (**A1**), pairs of noisy tiles are extracted (**A2**). The UNet model is trained using the mean-squared error between the model prediction and the other noisy frame (**A3**). **B1** Example of raw morphology frame. **B2** Frame denoised with the UNet model. **B3** Sharpened and enhanced frame with the deconvolution. **B4** Final segmentation of the ER structure.

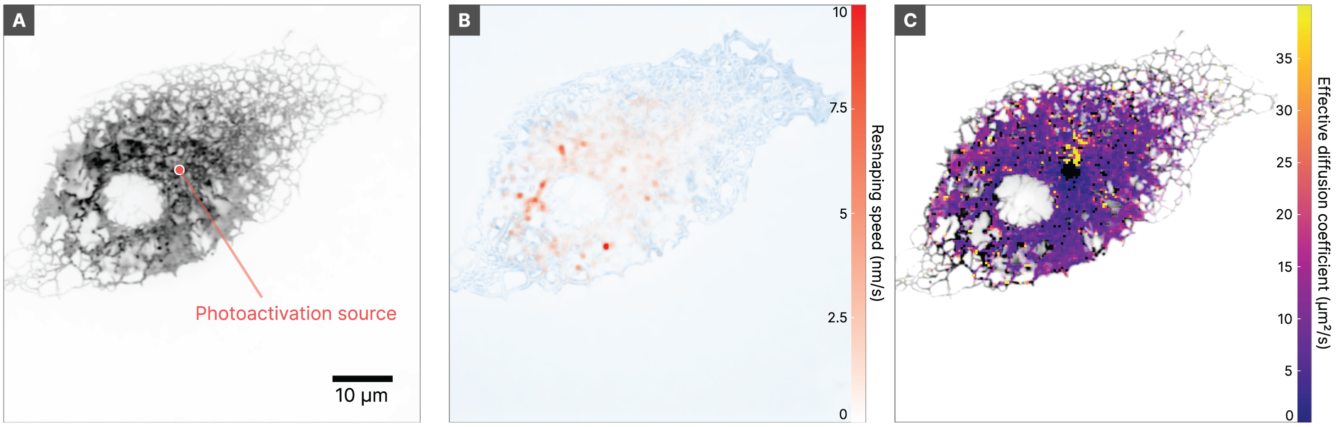


Figure S2. **Estimation of morphology reshaping. A** Cell morphology. **B** Reshaping intensity estimated by optical flow (red) and relative variance (blue shade). **C** Comparison with the diffusion map for the same cell.

| Parameter | Value |
|---|---|
| PSF $\sigma$ | 1.27 px |
| $\mu$ | 2 |
| $\lambda_1$ | 0.1 |
| $\lambda_v$ | 0.05 / N |
| $\lambda_t$ | 1 |
| Split Bregman inner iterations | 2 |
| Split Bregman outer iterations | 5 |

Table S1. **Parameter values for morphology deconvolution.** See equation 1

.

| Condition | N° of cells | N° of valid fits | Diffusion coefficient $D_{\text{eff}}$ | Effective MSD exponent $\alpha$ |
|---|---|---|---|---|
| HaloTag-KDEL | 39 | 34 | 16.84 (12.38, 24.70) μm$^2$ s$^{-1}$ | 0.997 (0.940, 1.048) |
| 3xHaloTag-KDEL | 17 | 17 | 8.53 (4.82, 10.78) μm$^2$ s$^{-1}$ | 1.008 (0.991, 1.029) |
| HaloTag-Sec61$\beta$ | 28 | 28 | 5.39 (4.04, 8.11) μm$^2$ s$^{-1}$ | 1.001 (0.834, 1.093) |
| 3xHaloTag-Sec61$\beta$ | 14 | 14 | 3.55 (2.79, 4.25) μm$^2$ s$^{-1}$ | 0.988 (0.925, 1.092) |
| CD3$\delta\Delta$-HaloTag | 50 | 47 | 6.19 (3.95, 8.41) μm$^2$ s$^{-1}$ | 1.005 (0.944, 1.113) |

Table S2. Median values (and 1st, 3rd quartiles) of diffusion coefficient $D_{\text{eff}}$ and effective MSD exponent ($\alpha$) of the different conditions. We also report the total number of cells analyzed and the number of successful fits.

| | $H$ statistic | $p$-value | Significance ($\alpha \leq 1 \times 10^{-3}$) |
|---|---|---|---|
| Difference in $D_{\text{eff}}$ between groups | 74.20 | $2.944 \times 10^{-15}$ | Yes |
| Difference in MSD exponent between groups | 1.73 | 0.785 | No |

Table S3. Kruskal–Wallis $H$ test to check whether there exists a significant difference in the distribution of either effective diffusion coefficient $D_{\text{eff}}$ or effective MSD exponent ($\alpha$).

| Group A | Group B | $U$ statistic | Test type | $p$-value (uncorrected) | $p$-value (corrected) | Significance ($\alpha \leq 1 \times 10^{-3}$) |
|---|---|---|---|---|---|---|
| HaloTag-KDEL | HaloTag-Sec61$\beta$ | 927.0 | one-sided (A > B) | $9.310 \times 10^{-11}$ | $2.793 \times 10^{-10}$ | Yes |
| HaloTag-KDEL | CD3$\delta\Delta$-HaloTag | 1505.0 | one-sided (A > B) | $7.322 \times 10^{-12}$ | $2.929 \times 10^{-11}$ | Yes |
| HaloTag-KDEL | 3xHaloTag-KDEL | 532.0 | one-sided (A > B) | $6.315 \times 10^{-7}$ | $1.263 \times 10^{-6}$ | Yes |
| HaloTag-Sec61$\beta$ | 3xHaloTag-Sec61$\beta$ | 290.0 | two-sided (A $\lesseqgtr$ B) | $1.261 \times 10^{-2}$ | $1.261 \times 10^{-2}$ | No |

Table S4. Pairwise Mann–Whitney–Wilcoxon $U$ test between conditions of interest. $p$-values were corrected for multiple comparisons via Holm–Bonferroni method.

#### A. Effective diffusion model

We consider a model of classical diffusion in a planar space. We assume that the recorded luminosity is proportional to the density of photoactivated particles in the given region. Let us call $\phi(t)$ the density of fluorescent particles in the source region $S$. Standard diffusion theory defines the density of photoactivated particles at any point in space by the following Cauchy problem:

$$\begin{cases} \partial_t u(\boldsymbol{x},t) - D\nabla^2_{\boldsymbol{x}} u(\boldsymbol{x},t) = 0, & \boldsymbol{x} \notin S \\ u(\boldsymbol{x},t) = \phi(t), & \boldsymbol{x} \in \partial S \\ u(\boldsymbol{x},t=0) = 0, & \end{cases} \tag{S1}$$

where we denoted the diffusion coefficient by $D$, we imposed $\phi(t)$ as Dirichlet boundary condition on the boundary of the source region $S$, and assumed a zero initial condition. By Duhamel's theorem, if $v$ is the solution of the auxiliary problem with a time independent boundary condition

$$\begin{cases} \partial_t v(\boldsymbol{x},t) - D\nabla^2_{\boldsymbol{x}} v(\boldsymbol{x},t) = 0, & \boldsymbol{x} \notin S \\ v(\boldsymbol{x},t) = 1, & \boldsymbol{x} \in \partial S \\ v(\boldsymbol{x},t=0) = 0, & \end{cases} \tag{S2}$$

then we can build a solution for eq. (S1) as a convolution between the auxiliary solution $v(r,t)$ and the time-varying boundary term:

$$u(\boldsymbol{x},t) = \int_0^t \phi(t-\tau)\, \partial_\tau v(\boldsymbol{x},\tau) \mathrm{d}\tau. \tag{S3}$$

We consider this problem in a 2D plane with radial symmetry where the source $S$ is a disk of radius $a$ and diffusion occurs in the space $r > a$. Expressing the Laplace operator in polar coordinates, the auxiliary Cauchy problem becomes:

$$\begin{cases} \partial_t v(r,t) - D\left(\partial_r^2 v(r,t) + \frac{1}{r}\partial_r v(r,t)\right) = 0, & r > a \\ v(r,t) = 1, & r = a \\ v(r,t=0) = 0. & \end{cases} \tag{S4}$$

By taking the Laplace transform of the diffusion equation, we obtain a modified Bessel equation of order 0:

$$z^2 \partial_z^2 \hat{v}(r,s) + z\partial_z \hat{v}(r,s) - z^2 \hat{v}(r,s) = 0, \qquad \text{with } z = \sqrt{s/D}\, r \tag{S5}$$

where $\hat{v}(r,s) = \mathscr{L}\{v(r,t)\}$ indicates the Laplace transform of $v(r,t)$. The only non-diverging solution of the modified Bessel equation is

$$\hat{v}(r,s) = A(s)\, K_0(\sqrt{s/D} r) \tag{S6}$$

where $K_0(z)$ is the modified Bessel function of the second kind (order 0) and $A(s)$ is a coefficient determined by imposing the boundary condition at $r = a$. The solution is thus

$$\hat{v}(r,s) = \frac{K_0(\sqrt{s/D}\, r)}{s\, K_0(\sqrt{s/D}\, a)}. \tag{S7}$$

To find the solution of the time-varying problem, we need to apply Duhamel's theorem (eq. S3):

$$u(r,t) = \int_0^t \phi(t-\tau)\ \left(\mathscr{L}^{-1}\left\{s\,\hat{v}(r,s)\right\}\big|_\tau\right) d\tau, \tag{S8}$$

where we used the differentiation property $\mathscr{L}\left\{\partial_t f\right\} = s\,\mathscr{L}\left\{f\right\}$.

### B. Supplemental Text and Discussion

#### 1. *Potential sources of anomalous scaling of mean squared displacements*

Extensive work in the literature has addressed the theoretical treatments of diffusion in various nonhomogenous media. These treatments largely have focused on the effects of retention by a structure or interaction compared to unrestricted diffusion. Appealingly, these have nearly uniformly concluded that interactions or confinements of any nature cause subdiffusive scaling of mean squared displacements, since diffusing particles are increasingly likely to encounter a source of restricted motion over longer time lags than shorter ones. Thus, our result that structural confinement in our system causes the appearance of superdiffusive scaling is quite surprising.

We present this observation here without a full theoretical understanding, which will be the focus of future work, but we propose one potential explanation. An inherent weakness of photoactivation compared to FRAP of large regions or single molecule tracking is that all of the molecules observed in the experiment start at essentially the same location in the structure. Thus, anisotropy in the density of paths from the photoactivation point or heterogeneity of molecular species with distinct diffusion rates will not be averaged out across the structure as they would be with many distinct starting points for molecules. We note there is a clear precident in the ER for both of these phenomena [S13, S16, S25], and we provide some examples of the latter in this paper (fig. 6). Future work will need to integrate multiple approaches using the same constructs and conditions to establish a quantitative framework for where these sources of apparent superdiffusion come from, but here we suffice it to say that we can effectively remove them with an appropriate normalization to the structure.

#### 2. *A note on the use of intensity-weighting as a proxy for connectivity*

Purely applying a proposed diffusion model to the distance through the structure will produce a non-linearly scaled distance transform, analogous to fig. 1H, which is not a good predictor of the heterogeneity in photoactivated content observed within the structure and relative arrival times. Consequently, in fig. 2, we invoked a central assumption utilized in most analysis of FRAP in complex structures, that fluorescence signal could be used as a proxy for connectivity [S22]. In practice, we achieve this by weighting the photoactivated channel according to the underlying moxGFP signal (fig. 2). Thus, regions with more ER in them that are likely to allow more paths for molecules to traverse will gain signal at a rate proportional to the amount of ER present as measured by the moxGFP fluorescence. The remarkably strong predictive power of this approach even in a highly heterogeneous cell suggests that this long-held assumption in many FRAP models could be largely accurate, supporting the significant success of the use of this tool in the endoplasmic reticulum.

However, we note that concerns about the universality of this assumption have been raised previously in the literature [S19, S20], and our recent work as well as that of others in mapping the nanoscale heterogeneity of the ER structure suggest that is possible to have very different connectivity within ER structures that appear similar at the resolution of confocal microscopy (reviewed in [S16]). Fortunately, our improved approach for mapping where signal exists using the moxGFP channel and the observed Fisher information can be applied spatially to directly test the degree to which this assumption breaks down (fig. 6). Although these regions are not frequent, they do exist with some cells within the population observed. It will be a subject of great interest in future work to identify the factors that control where this occurs and if it is associated with specific biological phenomena.

#### 3. *Comments on membrane diffusion and the presence or absence of locally cyclic boundary conditions*

An appealing aspect of early FRAP studies in the ER was the apparent consistency of extracted effective diffusion coefficients with the theoretically expected values for membrane diffusion, based on early membrane mixing studies and measurements of rotational diffusion as ascertained through spectroscopy [S4, S5, S18]. These classic studies, largely performed on the plasma membranes of specialized cells, yielded a range of effective diffusion coefficients that are consistent with the now well studied interactions of membrane proteins undergoing transient interactions with cytoskeleton (reviewed in [S3, S7, S10]).

However, it was clear even in early studies that the ER and Golgi membranes showed significantly elevated fluidity [S10], and that the apparent slowed diffusion observed was at least partially the result of structural confinement. Models based on the known ER structures at the time (tubules and sheets) predicted 3-fold increases in the "unwrapped" protein diffusion [S22], and estimations based upon the observed fractal dimension in peripheral tubules predicted as much as a 4-fold increase in effective diffusion coefficient [S19, S20]. We note that the resulting numbers from both of these studies are consistent with recent data from our group using several distinct structurally-informed single molecule tracking analyses to analyze one-dimensional diffusion along the visible axis of ER tubules in the periphery [S17, S23].

The numbers reported in this study for ensemble ER membrane diffusion (fig. 3) are higher by a factor of 2-3 from these previous estimates performed solely at the periphery of cells. At first glance, this may seem to be incongruous with our previous results, but upon closer examination we feel it is consistent. The ER in the periphery of tissue culture cells is very highly curved and exists almost completely in a tubular form [S6, S15, S16]. As such, these regions are best approximated by the models introduced in previous work, and the appearance of motion in these regions is heavily dominated by one dimension of membrane diffusion obeying cyclic boundary conditions (i.e. - Proteins diffusing laterally around ER tubules would have the appearance of remaining stationary in a two dimensional projection at the resolution of confocal microscopy as performed here).

The full-cell nature of the experiments in this paper necessitate that much of the ER analyzed is within the more dense perinuclear and intermediate regions of ER, where ER structures become much more convoluted and less well predicted by simple models (reviewed in [S16]). Many of these structures still maintain degrees of freedom for lateral diffusion that stretch across several pixels, and as such the full distance of a random walk on the surface may be visible in the 2D projection, creating a potential for elevation of the observed effective diffusion coefficient in the projected structure (by as much as 3-fold, based on the model of [S22]), a result consistent with the data presented here. Understanding the similarities and differences of protein and luminal diffusion in peripheral tubules compared to the dense, convoluted structures closer to the nucleus is an area of great interest for future work, since these domains of the ER have been suggested to facilitate distinct functions or have unique lipid compositions [S16]. This will require the development of technology for correlating nanoscale membrane topology to dynamic readouts of molecular redistribution in the network, however.

#### 4. *Stucturally-informed analysis of photoactivation data as an additional tool in the biophysicists' toolbox*

Several resources in the literature exist for comparing and contrasting the spatial and temporal scales of biophysical tools for assaying the dynamic nature of molecular components (see, for example [S10]), so we only briefly discuss here. In short, high temporal resolution approaches like FCS and high-speed single molecule tracking are less likely to lose very raid species in a population, but are more prone to artifacts as the result of sampling bias due to the relatively smaller spatial scales over which they can be realistically performed.

It is worth pointing out that when considering anomalous diffusion (here, we use the looser definition–generally nonlinear scaling of mean squared displacements with time), sources of nonlinearity can exist at several distinct spatial and temporal scales. As such, two techniques with non-overlapping windows may report transport phenomena characterized by sub- (or super-) diffusive scaling of mean squared displacements for entirely different reasons. Indeed, in the ER this appears to be nearly always the case for the literature we are aware of. For example, millisecond time scale FCS measurements suggest anomalous diffusion as a result of tracer engagement by protein folding machinery [S11], but not as a result of membrane shape (albeit in the absence of asymmetric cyclic boundary conditions) [S24]. However, single molecule tracking reveals strong contributions of membrane structure to anomalous diffusion over time scales of a few seconds [S23], and photoconversion of known misfolded proteins still is fit quite well with a simple diffusion model (see fig. 5). Thus, the true nature of a diffusive (or non-diffusive) process can only be understood completely by integrating multiple tools over distinct spatial and temporal scales to collectively create a complete image of the behavior of a molecule in a complex system.

Our approach using photoactivation provides some benefits in this space, allowing FRAP-scale kinetics over entire cells. We note that applying our first order structural correction to the ER, we are able to recover the effective diffusion coefficients that are consistent with work by FCS [S11, S13, S24] and with structural unwrapping based on the fractal dimesnions of the peripheral ER [S20], but we are also able to capture the effectively slowed diffusion caused by frequent rapid engagement of folding machinery that is historically challenging to capture with FRAP [S14]. However our approach is still limited by low signal to noise at longer distances (especially for slower proteins), and necessitates either local or radial averaging and the assumptions inherent in the selection of any particular diffusion model.

Perhaps the most promising component of this approach is due to its compatibility with faster approaches like FCS and single molecule tracking. Historically, integrations of these tools across several scales has been effective in identifying distinct molecular states [S13] and effects of geometry on transport phenomena [S1], both factors likely to be at play in the ER. Future work will attempt to use structurally-informed photoactivation analysis in combination with FRAP, FLIP, FCS, and single molecule techniques to understand the contributions of local heterogeneity to the ensemble phenomena we have been able to describe here.

#### 5. *Discussion of implications and caveats of analyzing misfolded protein motion in the ER*

In this section, we note a few interesting points that arise from the analysis of the motion of the misfolded protein construct CD3$\delta\Delta$–HaloTag in the ER. To begin, recall that our approach comes with two inherent weaknesses in analyzing the motion of a protein with known multiple states. First, we assume the data to hold a single effective diffusion coefficient when performing the model fit. In the context of a misfolded protein that engages with folding machinery, this is almost certainly not the case. (Unfortunately, it has been well established that distinguishing two populations with different effective diffusion coefficients is very challenging or impossible unless the ratio of their $D_{\text{eff}}$s is very far from one, see [S1] for a nice example of this, so we must suffice here to detect decreases in the average $D_{\text{eff}}$ rather than direct resolution of two states).Second, the interference of scattered light from the photoactivation laser makes the model fit very poor in immediate vicinity of the photoactivation point, so analysis of a resistant fraction is challenging.

Taken together, these preclude the use of our data to evaluate the presence of a fraction of CD3$\delta\Delta$–HaloTag molecules that are completely immobilized. Those at the photoactivation point cannot be analyzed, and those away from the photoactivation point will not have ever been photoactivated and as such remain dark. However, this provides a benefit, since all the signal observed was by definition mobile at some point, so reductions in the effective diffusion coefficient are the result of interactions of mobile particles with folding machinery. Indeed, we observe exactly that. The effective diffusion coefficient of CD3$\delta\Delta$–HaloTag is lower than a luminal construct more than double the size, 3xHaloTag-KDEL. However, we note that in the absence of structural correction, the CD3$\delta\Delta$–HaloTag MSD scales even more superdiffusively than the much faster HaloTag-KDEL, and while structural correction returns the value of the scaling exponent to one, there are a few cells that still show mildly superdiffusive scaling after correction (fig. 5D, right tail). Since this construct is known to be misfolded and has been clearly demonstrated via biochemistry to be engaging the misfolded protein response machinery [S2, S12], we feel this highlights the risk of multiple distinct species to create the appearance of superdiffusive motion in photoactivation experiments. Data collected from parts of the ER that are far away from the photoactivation point will be expected to be biased towards larger fractions of molecules that are in more mobile states (see *Supplementary Text and Discussion*, Section 1 for full discussion).

In addition, we note that the inability of our approach to directly observe bound states of the molecule as subdiffusive scaling of the MSD exponent has three potential explanations. First, the size and temporal scale of this anomality may be beneath our temporal or spatial resolution. Thus far, engagement of misfolded cargo by folding machinery has only been reported by FCS [S11] as opposed to FRAP [S14]. We note that in contrast to [S14], we are able to see reduction of the effective diffusion coefficient in our data, though this is likely in part because we are working with a soluble cargo, where binding by membrane-associated machinery has a more dramatic reduction in the transport phenomena than for the membrane-bound VSV-G used in the previous studies. We note that this is in agreement with the well-established capacity of FRAP to visualize immobilization of soluble folding factors upon expression of a misfolded target [S8, S9, S21]. Second, as discussed above and in *Supplementary Text and Discussion*, Section 1, the presence of functionally distinct species within the molecular population can create the (false) appearance of superdiffusive scaling, and this may be dominating over the effects of subdiffusive trapping. Last, it worth considering that the subdiffusive scaling of MSDs may not be detectable if the complex of CD3$\delta\Delta$–HaloTag is still diffusing, just more slowly. This is exacerbated if the majority of the CD3$\delta\Delta$–HaloTag is in this bound state, which is completely plausible given the slow effective diffusion coefficient reported (fig. 5). If this is the case, it suggests that the protein folding or misfolded protein recognition machinery involved is quite dynamic in its own right, which will be an interesting subject for future research.

#### 6. *Implications of uneven time of arrival within the ER*

Our observation that exploration times in the ER are not always easily predictable or uniform without direct observation is interesting, since this uneven mixing could facilitate the development of spatial variation in the execution of ER-associated functions–a long standing puzzle in the field of ER biology. If this is the case, one would predict that this capacity must be able to be directly regulated in a location-specific manner. We suggest here one way this could potentially be locally controlled, through the direct modulation of ER structure at the nanoscale via either external forces or the activity of local ER-shaping proteins.

Correlative work with single molecule tracking, photoactivation, and simulations has been performed in the cytoplasm of E. coli, which have a highly reproducible cylindrical shape [S1]. A major conclusion of the work was that the mixing speed was dependent on both the length that must be traveled between two points along the central cylindrical axis and the radius of the cylinder, since this latter variable significantly affects the density and distribution of potential paths away from the location of the molecule at the initial time point. Earlier work removing the contribution of the ER tubule structure in single molecule trajectories accounted for the length, but largely ignored the contributions of the radial dimensions, since in the ER of mammalian cells these are well beneath the resolution limit of conventional microscopy [S23]. However, it stands to reason that the surface area of a slice through the longitudinal axis of any particular ER structure may be a significant contributor to the rate of flux through the structure, a variable we do not address in this work.

Early work on the structure of the ER largely viewed the system as a network of two relatively simple shapes, sheets and tubules. In the era of three dimensional electron microscopy, this view has been challenged by numerous intermediate forms and more complex shapes at the nanoscale [S16]. Unfortunately, these shapes have highly variable membrane-to-lumen ratios, and as such are very difficult to predict their effects on transport. Thus, in this work we have loosely depended on the observed fluorescence as a proxy for connectivity (see *Supplementary Text and Discussion*, Section 2), but we note that our ability to predict this accurately decreases as the ER becomes more complex in the perinuclear and transitional regions close to the nucleus. We note that most regions of more rapid arrival time are in this dense area, and we have previously demonstrated that the frequency of ER luminal content increases moving into this area [S25]. Thus, these regions may represent places where elevated connectivity or decreased surface area-to-volume ratios facilitate more rapid mixing of luminal content. This is consistent with numerous observations showing elevated rates of specific ER functions within this region (reviewed in [S16]), an interesting direction for future work.

---


[S1] Somenath Bakshi, Benjamin P Bratton, and James C Weisshaar. Subdiffraction-limit study of kaede diffusion and spatial distribution in live escherichia coli. *Biophysical journal*, 101(10):2535–2544, 2011.

[S2] Riccardo Bernasconi, Carmela Galli, Verena Calanca, Toshihiro Nakajima, and Maurizio Molinari. Stringent requirement for hrd1, sel1l, and os-9/xtp3-b for disposal of erad-ls substrates. *Journal of Cell Biology*, 188(2):223–235, 2010.

[S3] Michael Edidin. Patches and fences: probing for plasma membrane domains. *Journal of Cell Science*, 1993(Supplement_17):165–169, 1993.

[S4] Michael Edidin and TY Wei. Diffusion rates of cell surface antigens of mouse-human heterokaryons. i. analysis of the population. *The Journal of cell biology*, 75(2):475–482, 1977.

[S5] Larry D Frye and Michael Edidin. The rapid intermixing of cell surface antigens after formation of mouse-human heterokaryons. *Journal of cell science*, 7(2):319–335, 1970.

[S6] Larissa Heinrich, Davis Bennett, David Ackerman, Woohyun Park, John Bogovic, Nils Eckstein, Alyson Petruncio, Jody Clements, Song Pang, C Shan Xu, et al. Whole-cell organelle segmentation in volume electron microscopy. *Nature*, 599(7883):141–146, 2021.

[S7] Darius V Köster and Satyajit Mayor. Cortical actin and the plasma membrane: inextricably intertwined. *Current opinion in cell biology*, 38:81–89, 2016.

[S8] Chun Wei Lai, Deborah E Aronson, and Erik Lee Snapp. Bip availability distinguishes states of homeostasis and stress in the endoplasmic reticulum of living cells. *Molecular biology of the cell*, 21(12):1909–1921, 2010.

[S9] Chunwei Walter Lai, Joel H Otero, Linda M Hendershot, and Erik Snapp. Erdj4 protein is a soluble endoplasmic reticulum (er) dnaj family protein that interacts with er-associated degradation machinery. *Journal of Biological Chemistry*, 287(11):7969–7978, 2012.

[S10] Jennifer Lippincott-Schwartz, Erik Snapp, and Anne Kenworthy. Studying protein dynamics in living cells. *Nature reviews Molecular cell biology*, 2(6):444–456, 2001.

[S11] Nina Malchus and Matthias Weiss. Anomalous diffusion reports on the interaction of misfolded proteins with the quality control machinery in the endoplasmic reticulum. *Biophysical journal*, 99(4):1321–1328, 2010.

[S12] Martin Mehnert, Thomas Sommer, and Ernst Jarosch. Erad ubiquitin ligases: multifunctional tools for protein quality control and waste disposal in the endoplasmic reticulum. *Bioessays*, 32(10):905–913, 2010.

[S13] Hisao Nagaya, Taku Tamura, Arisa Higa-Nishiyama, Koji Ohashi, Mayumi Takeuchi, Hitoshi Hashimoto, Kiyotaka Hatsuzawa, Masataka Kinjo, Tatsuya Okada, and Ikuo Wada. Regulated motion of glycoproteins revealed by direct visualization of a single cargo in the endoplasmic reticulum. *The Journal of Cell Biology*, 180(1):129–143, 2008.

[S14] Sarah Nehls, Erik L Snapp, Nelson B Cole, Kristien JM Zaal, Anne K Kenworthy, Theresa H Roberts, Jan Ellenberg, John F Presley, Eric Siggia, and Jennifer Lippincott-Schwartz. Dynamics and retention of misfolded proteins in native er membranes. *Nature cell biology*, 2(5):288–295, 2000.

[S15] Jonathon Nixon-Abell, Christopher J Obara, Aubrey V Weigel, Dong Li, Wesley R Legant, C Shan Xu, H Amalia Pasolli, Kirsten Harvey, Harald F Hess, Eric Betzig, et al. Increased spatiotemporal resolution reveals highly dynamic dense tubular matrices in the peripheral ER. 354(6311):aaf3928.

[S16] Christopher J Obara, Andrew S Moore, and Jennifer Lippincott-Schwartz. Structural diversity within the endoplasmic reticulum—from the microscale to the nanoscale. *Cold Spring Harbor Perspectives in Biology*, page a041259, 2022.

[S17] Christopher J Obara, Jonathon Nixon-Abell, Andrew S Moore, Federica Riccio, David P Hoffman, Gleb Shtengel, C Shan Xu, Kathy Schaefer, H Amalia Pasolli, Jean-Baptiste Masson, et al. Motion of single molecular tethers reveals dynamic subdomains at er-mitochondria contact sites. *bioRxiv*, pages 2022–09, 2022.

[S18] Mu-Ming Poo and Richard A Cone. Lateral diffusion of rhodopsin in the photoreceptor membrane. *Nature*, 247(5441):438–441, 1974.

[S19] Ivo F Sbalzarini, Arnold Hayer, Ari Helenius, and Petros Koumoutsakos. Simulations of (an) isotropic diffusion on curved biological surfaces. *Biophysical journal*, 90(3):878–885, 2006.

[S20] Ivo F Sbalzarini, Anna Mezzacasa, Ari Helenius, and Petros Koumoutsakos. Effects of organelle shape on fluorescence recovery after photobleaching. *Biophysical journal*, 89(3):1482–1492, 2005.

[S21] Jingshi Shen, Erik L Snapp, Jennifer Lippincott-Schwartz, and Ron Prywes. Stable binding of atf6 to bip in the endoplasmic reticulum stress response. *Molecular and cellular biology*, 25(3):921–932, 2005.

[S22] Eric D Siggia, Jennifer Lippincott-Schwartz, and Stefan Bekiranov. Diffusion in inhomogeneous media: theory and simulations applied to whole cell photobleach recovery. *Biophysical journal*, 79(4):1761–1770, 2000.

[S23] Yunhao Sun, Zexi Yu, Christopher J. Obara, Keshav Mittal, Jennifer Lippincott-Schwarz, and Elena F. Koslover. Unraveling Single-Particle Trajectories Confined in Tubular Networks.

[S24] Matthias Weiss, Hitoshi Hashimoto, and Tommy Nilsson. Anomalous protein diffusion in living cells as seen by fluorescence correlation spectroscopy. *Biophysical journal*, 84(6):4043–4052, 2003.

[S25] Pengli Zheng, Christopher J Obara, Ewa Szczesna, Jonathon Nixon-Abell, Kishore K Mahalingan, Antonina Roll-Mecak, Jennifer Lippincott-Schwartz, and Craig Blackstone. Er proteins decipher the tubulin code to regulate organelle distribution. *Nature*, 601(7891):132–138, 2022.
